# A Novel Glycoproteomics Platform for High-Throughput Identification of Disease-Associated Glycoforms

**DOI:** 10.64898/2026.03.06.708406

**Authors:** Shengye Wen, Yu Gao, Xinyu Miao, Jianbo Deng, Yufeng Zhou, Wei Ge, Siyue Bo, Wenqi Zhang, Rumeng Zhang, Chunyan Hou, Junfeng Ma, Junhong Jiang, Shuang Yang

**Affiliations:** Laboratory of Clinical and Molecular Glycobiology, Institute of Glycomics, Shantou University Medical College, Shantou, Guangdong 515041, China; Clinical Medicine Research Center, The First Affiliated Hospital of Shantou University Medical College, Shantou, Guangdong 515041, China; Center for Clinical Mass Spectrometry, School of Pharmaceutical Sciences, Soochow University, Jiangsu 215123, China; Department of Respiratory Medicine, The Fourth Affiliated Hospital of Soochow University, Suzhou, Jiangsu 215123, China; Department of Oncology, Lombardi Comprehensive Cancer Center, Georgetown University Medical Center, Georgetown University, Washington, DC 20057, USA

**Author notes:** All inquiries correspondence to S.Y., Shantou University Medical College. These authors contributed equally to this work (Shengye Wen, Yu Gao, Xinyu Miao, Jianbo Deng, Yufeng Zhou).

**Keywords:** Biomarker, Glycosylation, Glycoenzyme, Glycopattern, Neurodegenerative disease, Mass spectrometry

## Abstract

Glycosylation is a critical post-translational modification, and its aberrant forms are potent disease biomarkers; however, the comprehensive, site-specific identification of all glycosites and glycoforms across an entire proteome remains prohibitively slow and computationally demanding. To address this bottleneck and accelerate biomarker discovery, we introduce the **Glycoproteomics Data Analysis Software** (GDAS), a novel, high-throughput platform designed to provide confident, proteome-scale identification of disease-specific glycoforms. GDAS streamlines the analysis through a core, multi-step workflow: it initially employs an ultrafast open search (e.g., MSFragger-Glyco) on mass spectrometry data to rapidly screen and statistically reduce the vast proteome database to a manageable subset of significantly regulated glycoproteins, conserving computational resources for subsequent, in-depth, targeted N- and O-glycosylation analysis using specialized tools (e.g., GlycReSoft and O-Pair). Furthermore, a unique Final Analysis Module utilizes an advanced statistical and machine learning pipeline (incorporating Bootstrap/Bayesian methods, XGBoost, and Random Forest) to integrate quantitative results and generate a robust, comprehensive glycosylation score. We demonstrate GDAS’s power to recognize biologically relevant glycosylation changes in targeted proteins by validating it using published Alzheimer’s disease data. GDAS can be downloaded from https://github.com/yang-lab/GDAS.

**Graphical Abstract:** 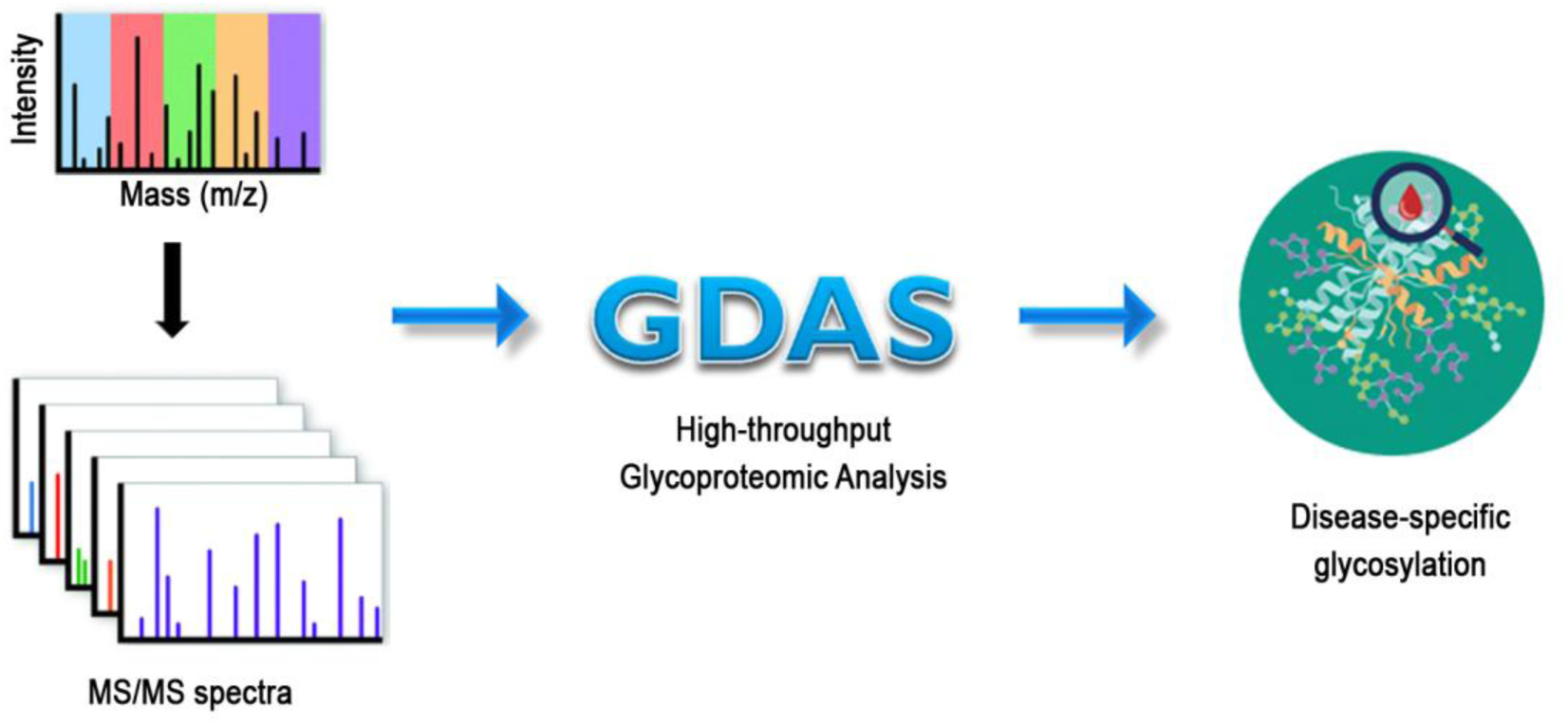

## Introduction

Glycosylation is a continuously expanding field as it occurs not only on protein or lipid but also on ribonucleic acid (RNA) ^1,2^. These modifications are crucial for maintaining healthy cell function, while disturbances in glycosylation can lead to the occurrence and progression of diseases. Defective activity of glycoenzymes is found in inborn errors of metabolism known as congenital disorders of glycosylation (CDG), with phosphomannomutase 2-CDG (PMM2-CDG or CDG-1a) being the most common example ^3,4^. Other less common autosomal dominant inheritance patterns may involve defective activity in O-linked glycosylation, such as with exostosin-1 (EXT1) or EXT2 ^5^, and POFUT1 or POGLUT1 ^6^. In cancer, it is widely acknowledged that aberrant glycosylation can define tumor malignancy, meaning that tumor cells have unique characteristics in their glycopatterns, including both their glycan profile and site-specific glycosylation ^7^. Cancer cells generally show higher expression of immature truncated O-glycans, branched N-glycans, and core-fucosylated N-glycans ^8–10^. These changes are a result of dysregulated glycoenzymes in the tumor microenvironment (TME), which leads to different glycan synthesis on their substrates ^11^. Altered glycosylation has been implicated in cell adhesion and immune recognition and, quite importantly, is used for cancer biomarkers and therapeutics ^12^.

Mass spectrometry (MS) has long been a powerful technique for characterizing complex glycopatterns by analyzing intact glycoconjugates (precursor ions) and their fragment ions ^13^. The analysis typically uses a multi-pronged approach, combining various ionization and fragmentation techniques to fully characterize the glycoprotein structure, including the glycan composition (N-glycans or O-glycans) and site of attachment on the protein (asparagine = N, serine = S, threonine = T) ^14^. Initially, intact glycopeptides are analyzed using soft ionization techniques like electrospray ionization (ESI) to determine their molecular weight ^15^. Matrix-assisted laser desorption/ionization (MALDI) is commonly used to determine molecular weight and, with necessary modifications, may even provide information on glycan compositions and structure ^16–18^. For more structural detail, fragmentation techniques are used to target peptides and glycans. Collision-induced dissociation (CID) and higher-energy collisional dissociation (HCD) are commonly used to generate b-, y-, c- or z-ions from glycopeptides, but they often preferentially break the weaker glycosidic bonds ^19^. This can be useful for sequencing the glycan but makes localizing the exact attachment site challenging, especially for site-specific O-glycosylation assignment ^20^. In contrast, electron transfer dissociation (ETD, predominantly c-/z-ions) and electron-transfer/higher-energy collision dissociation (EThcD, b-/y-/c-/z-ions and oxonium ions) offers a distinct advantage by preferentially cleaving the peptide backbone while leaving the labile glycosidic bonds intact ^21^. This is ideal for localizing the glycan attachment site and sequencing the peptide. Furthermore, ion mobility spectrometry (IMS) can be coupled with MS to separate glycopeptides based on their size, shape, and charge, helping to resolve isomers and enhance structural characterization ^22^. By integrating these various ionization and fragmentation techniques, researchers can now obtain a comprehensive fingerprint of complex glycosylation, from the overall glycoprotein mass to the fine details of the glycan structure and its specific location.

The analysis of MS data for protein glycosylation relies on several computational approaches, primarily categorized by their search logic: peptide-first, glycan-first, or hybrid search engines. These algorithms are either implemented by matching MS data to known protein and/or glycan databases (database-driven (DD)) ^23^ or by inferring the sequence from scratch (de novo (DN)) ^24^. For instance, GlycoMod can predict the composition of intact glycopeptides from MS1 data by using a list of experimental masses, adduct ions, and monosaccharide residues ^25^. Similarly, SimGlycan predicts glycan structures from MS2 data and is also capable of high-throughput analysis of intact glycopeptides ^15^. Since their development, numerous software platforms have been established for studying both N- and O-glycopeptides (**Table 1**).

**Table 1.**
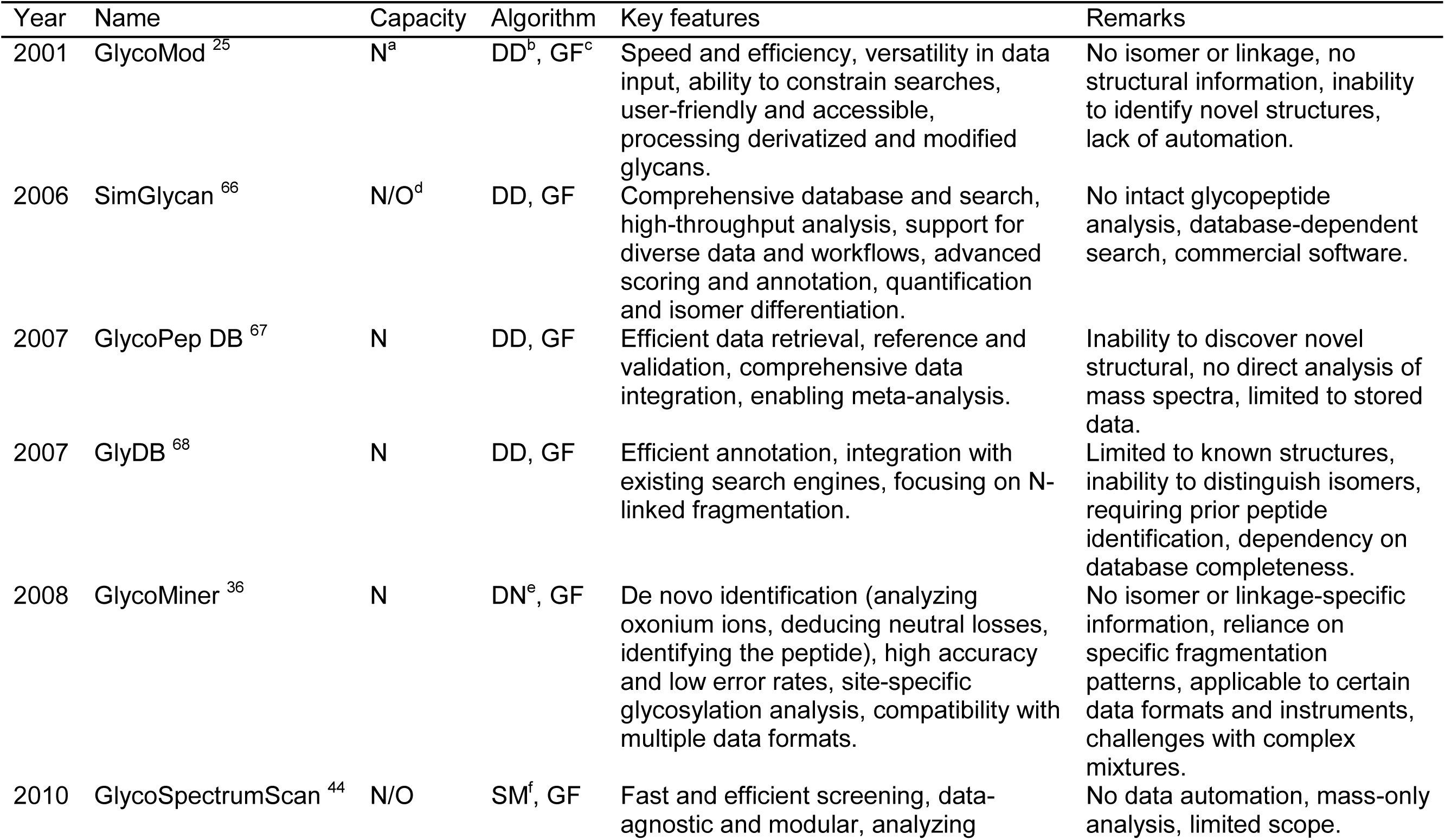
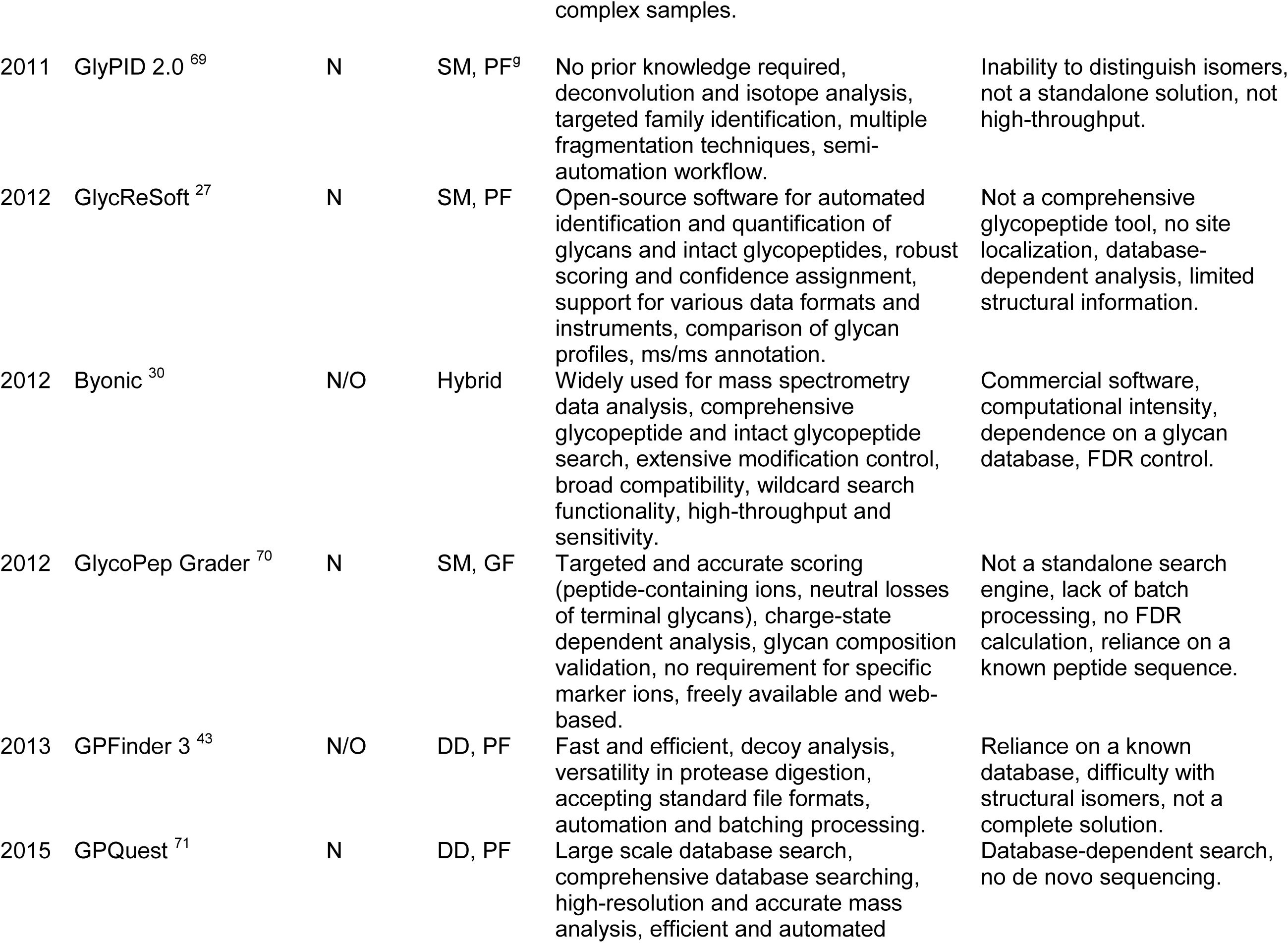
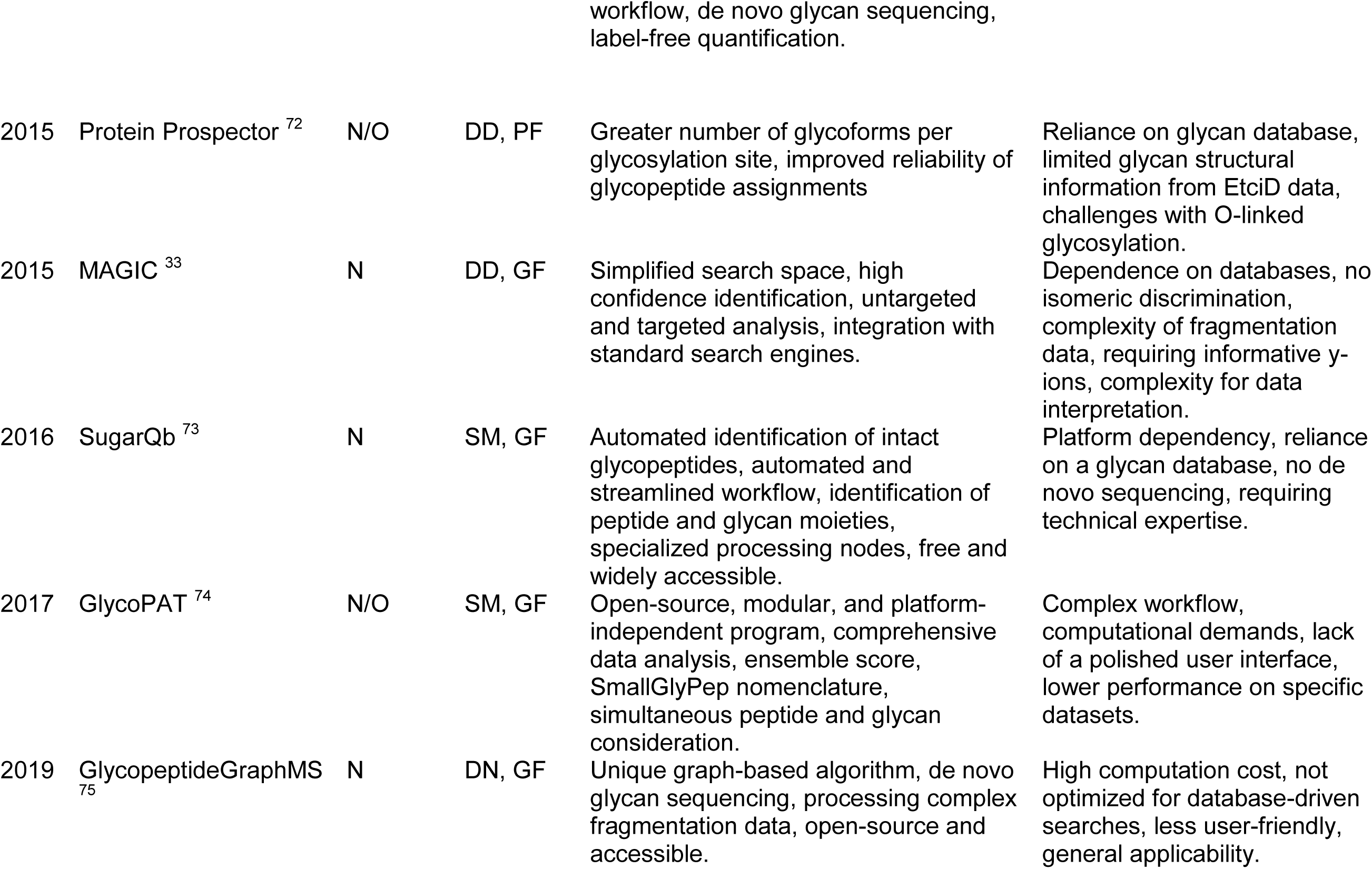
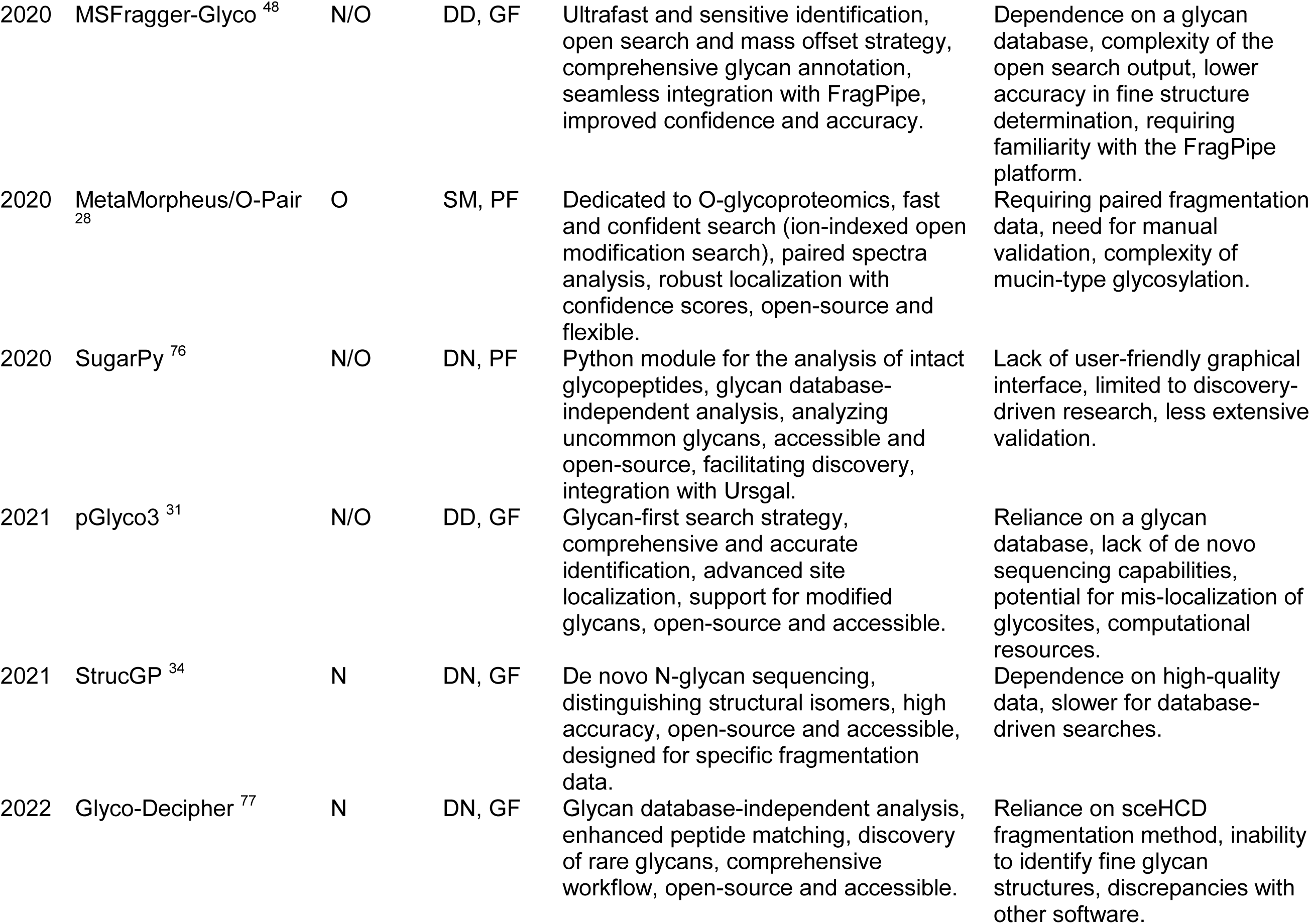
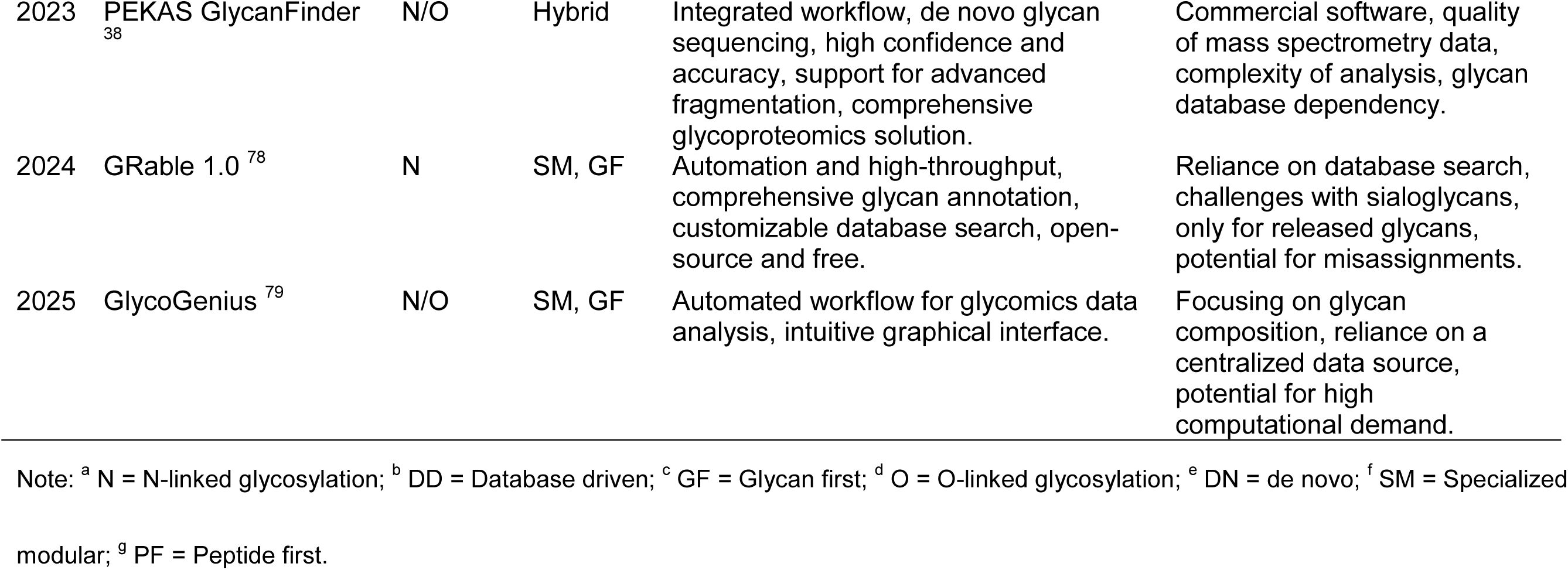
Mass spectrometry software for glycosylation analysis. This table lists various software tools used for analyzing N-linked and/or O-linked glycosylation via mass spectrometry. It includes a brief overview of each software’s key features; more detailed information can be found in their respective publications.

| Year | Name | Capacity | Algorithm | Key features | Remarks |
| --- | --- | --- | --- | --- | --- |
| 2001 | GlycoMod <sup>25</sup> | N<sup>a</sup> | DD<sup>b</sup>, GF<sup>c</sup> | Speed and efficiency, versatility in data input, ability to constrain searches, user-friendly and accessible, processing derivatized and modified glycans. | No isomer or linkage, no structural information, inability to identify novel structures, lack of automation. |
| 2006 | SimGlycan <sup>66</sup> | N/O<sup>d</sup> | DD, GF | Comprehensive database and search, high-throughput analysis, support for diverse data and workflows, advanced scoring and annotation, quantification and isomer differentiation. | No intact glycopeptide analysis, database-dependent search, commercial software. |
| 2007 | GlycoPep DB <sup>67</sup> | N | DD, GF | Efficient data retrieval, reference and validation, comprehensive data integration, enabling meta-analysis. | Inability to discover novel structural, no direct analysis of mass spectra, limited to stored data. |
| 2007 | GlyDB <sup>68</sup> | N | DD, GF | Efficient annotation, integration with existing search engines, focusing on N-linked fragmentation. | Limited to known structures, inability to distinguish isomers, requiring prior peptide identification, dependency on database completeness. |
| 2008 | GlycoMiner <sup>36</sup> | N | DN<sup>e</sup>, GF | De novo identification (analyzing oxonium ions, deducing neutral losses, identifying the peptide), high accuracy and low error rates, site-specific glycosylation analysis, compatibility with multiple data formats. | No isomer or linkage-specific information, reliance on specific fragmentation patterns, applicable to certain data formats and instruments, challenges with complex mixtures. |
| 2010 | GlycoSpectrumScan <sup>44</sup> | N/O | SM<sup>f</sup>, GF | Fast and efficient screening, data-agnostic and modular, analyzing | No data automation, mass-only analysis, limited scope. |
| 2011 | GlyPID 2.0 <sup>69</sup> | N | SM, PF<sup>9</sup> | No prior knowledge required, deconvolution and isotope analysis, targeted family identification, multiple fragmentation techniques, semi-automation workflow. | Inability to distinguish isomers, not a standalone solution, not high-throughput. |
| 2012 | GlycReSoft <sup>27</sup> | N | SM, PF | Open-source software for automated identification and quantification of glycans and intact glycopeptides, robust scoring and confidence assignment, support for various data formats and instruments, comparison of glycan profiles, ms/ms annotation. | Not a comprehensive glycopeptide tool, no site localization, database-dependent analysis, limited structural information. |
| 2012 | Byonic <sup>30</sup> | N/O | Hybrid | Widely used for mass spectrometry data analysis, comprehensive glycopeptide and intact glycopeptide search, extensive modification control, broad compatibility, wildcard search functionality, high-throughput and sensitivity. | Commercial software, computational intensity, dependence on a glycan database, FDR control. |
| 2012 | GlycoPep Grader <sup>70</sup> | N | SM, GF | Targeted and accurate scoring (peptide-containing ions, neutral losses of terminal glycans), charge-state dependent analysis, glycan composition validation, no requirement for specific marker ions, freely available and web-based. | Not a standalone search engine, lack of batch processing, no FDR calculation, reliance on a known peptide sequence. |
| 2013 | GPFinder 3 <sup>43</sup> | N/O | DD, PF | Fast and efficient, decoy analysis, versatility in protease digestion, accepting standard file formats, automation and batching processing. | Reliance on a known database, difficulty with structural isomers, not a complete solution. |
| 2015 | GPQuest <sup>71</sup> | N | DD, PF | Large scale database search, comprehensive database searching, high-resolution and accurate mass analysis, efficient and automated | Database-dependent search, no de novo sequencing. |
| 2015 | Protein Prospector <sup>72</sup> | N/O | DD, PF | Greater number of glycoforms per glycosylation site, improved reliability of glycopeptide assignments | Reliance on glycan database, limited glycan structural information from EtcID data, challenges with O-linked glycosylation. |
| 2015 | MAGIC <sup>33</sup> | N | DD, GF | Simplified search space, high confidence identification, untargeted and targeted analysis, integration with standard search engines. | Dependence on databases, no isomeric discrimination, complexity of fragmentation data, requiring informative y-ions, complexity for data interpretation. |
| 2016 | SugarQb <sup>73</sup> | N | SM, GF | Automated identification of intact glycopeptides, automated and streamlined workflow, identification of peptide and glycan moieties, specialized processing nodes, free and widely accessible. | Platform dependency, reliance on a glycan database, no de novo sequencing, requiring technical expertise. |
| 2017 | GlycoPAT <sup>74</sup> | N/O | SM, GF | Open-source, modular, and platform-independent program, comprehensive data analysis, ensemble score, SmallGlyPep nomenclature, simultaneous peptide and glycan consideration. | Complex workflow, computational demands, lack of a polished user interface, lower performance on specific datasets. |
| 2019 | GlycopeptideGraphMS <sup>75</sup> | N | DN, GF | Unique graph-based algorithm, de novo glycan sequencing, processing complex fragmentation data, open-source and accessible. | High computation cost, not optimized for database-driven searches, less user-friendly, general applicability. |
| 2020 | MSFragger-Glyco<sup>48</sup> | N/O | DD, GF | Ultrafast and sensitive identification, open search and mass offset strategy, comprehensive glycan annotation, seamless integration with FragPipe, improved confidence and accuracy. | Dependence on a glycan database, complexity of the open search output, lower accuracy in fine structure determination, requiring familiarity with the FragPipe platform. |
| 2020 | MetaMorpheus/O-Pair<sup>28</sup> | O | SM, PF | Dedicated to O-glycoproteomics, fast and confident search (ion-indexed open modification search), paired spectra analysis, robust localization with confidence scores, open-source and flexible. | Requiring paired fragmentation data, need for manual validation, complexity of mucin-type glycosylation. |
| 2020 | SugarPy<sup>76</sup> | N/O | DN, PF | Python module for the analysis of intact glycopeptides, glycan database-independent analysis, analyzing uncommon glycans, accessible and open-source, facilitating discovery, integration with Ursgal. | Lack of user-friendly graphical interface, limited to discovery-driven research, less extensive validation. |
| 2021 | pGlyco3<sup>31</sup> | N/O | DD, GF | Glycan-first search strategy, comprehensive and accurate identification, advanced site localization, support for modified glycans, open-source and accessible. | Reliance on a glycan database, lack of de novo sequencing capabilities, potential for mis-localization of glycosites, computational resources. |
| 2021 | StrucGP<sup>34</sup> | N | DN, GF | De novo N-glycan sequencing, distinguishing structural isomers, high accuracy, open-source and accessible, designed for specific fragmentation data. | Dependence on high-quality data, slower for database-driven searches. |
| 2022 | Glyco-Decipher<sup>77</sup> | N | DN, GF | Glycan database-independent analysis, enhanced peptide matching, discovery of rare glycans, comprehensive workflow, open-source and accessible. | Reliance on sceHCD fragmentation method, inability to identify fine glycan structures, discrepancies with other software. |
| 2023 | PEKAS GlycanFinder<sup>38</sup> | N/O | Hybrid | Integrated workflow, de novo glycan sequencing, high confidence and accuracy, support for advanced fragmentation, comprehensive glycoproteomics solution. | Commercial software, quality of mass spectrometry data, complexity of analysis, glycan database dependency. |
| 2024 | GRable 1.0<sup>78</sup> | N | SM, GF | Automation and high-throughput, comprehensive glycan annotation, customizable database search, open-source and free. | Reliance on database search, challenges with sialoglycans, only for released glycans, potential for misassignments. |
| 2025 | GlycoGenius<sup>79</sup> | N/O | SM, GF | Automated workflow for glycomics data analysis, intuitive graphical interface. | Focusing on glycan composition, reliance on a centralized data source, potential for high computational demand. |

Note: <sup>a</sup> N = N-linked glycosylation; <sup>b</sup> DD = Database driven; <sup>c</sup> GF = Glycan first; <sup>d</sup> O = O-linked glycosylation; <sup>e</sup> DN = de novo; <sup>f</sup> SM = Specialized modular; <sup>g</sup> PF = Peptide first.

Peptide-first approaches are based on the premise that the search engine first identifies the peptide backbone, treating the glycan as a variable modification with a unique combined mass^26^. This strategy is widely used for intact glycopeptide analysis in software like GlycReSoft ^27^, MetaMorpheus/O-Pair ^28^, FragPipe/MSFragger ^29^, and Byonic ^30^. In contrast, the glycan-first strategy, exemplified by pGlyco3 ^31^, prioritizes identifying the glycan structure before analyzing the peptide backbone, which allows for a more comprehensive analysis of complex glycan modifications. Notably, some platforms like MetaMorpheus and FragPipe also offer glycan-first options to identify glycans prior to peptide matching ^32^. A related strategy, known as glycan-removal, identifies the peptide part of a glycopeptide based on specific peptide ions before analyzing the glycan structure. Examples of this include MAGIC ^33^, StrucGP ^34^, and Glyco-Decipher ^35^, all of which are effective for identifying intact glycopeptides across various MS fragmentation modes.

Several specialized software tools have been developed to tackle the complex task of identifying site-specific N-glycosylation, which involves linking a specific glycan to its exact amino acid location on a peptide. Early on, GlycoMiner ^36^ used a de novo approach to perform this analysis by examining fragmentation ions, while GlycReSoft ^27^ supported intact glycopeptide analysis but was limited in its ability to pinpoint the exact site. Byonic ^30^ emerged as a leading commercial option, excelling in accurate, site-specific searches using comprehensive databases and advanced scoring. However, Byonic has not yet been used to search a large protein database for complex protein glycosylation, such as N-glycosylation in the entire human proteome. More recent tools like pGlyco3 ^31^ have further refined this capability with advanced algorithms that combine different types of mass spectra for confident site localization. Additionally, platforms like Protein Prospector ^37^ and PEAKS Glycopeakfinder ^38^ also incorporate features for identifying site-specific glycoforms and resolving structural ambiguities, often by combining both database and de novo search strategies.

A primary challenge in glycosylation analysis is the complexity and heterogeneity of both O-glycans and their attachment sites. Unlike N-glycosylation, which is limited to asparagine, O-glycosylation can occur on a broader range of residues, including S, T, tyrosine (Y), hydroxylysine, and hydroxyproline ^39^. While most software focuses on N-glycosylation, a growing number of tools are specifically designed to handle the complexities of site-specific O-glycosylation. As highlighted in **Table 1**, hybrid platforms like Byonic and pGlyco3 can analyze both N- and O-linked glycosylation, offering comprehensive searches and advanced site localization. More specialized tools have also emerged, such as MetaMorpheus/O-Pair, an open-source platform dedicated to O-glycoproteomics that uses a specialized ion-indexed search strategy and paired fragmentation data for robust localization with confidence scores ^28,40^. Additionally, PEKAS GlycanFinder and Protein Prospector are recognized for their ability to analyze O-linked glycosylation and assign glycoforms to specific sites ^41^, demonstrating the increasing development of accurate tools for this challenging area of proteomics. However, searching an entire proteome database (e.g., human, mouse) for protein glycosylation, especially O-glycosylation, is time-consuming and computationally intensive, which can lead to software crashes ^42^.

To overcome the inherent computational challenges of glycoproteomics, a sophisticated, multi-step workflow is often employed. Several software tools are known for their rapid analysis of intact glycopeptides. MSFragger-Glyco uses an open search and mass offset strategy for ultrafast and sensitive identification ^28^. Similarly, GPFinder 3 is noted for its speed and efficiency, supporting automation and batch processing ^43^, while GlycoSpectrumScan offers a fast and efficient screening of intact glycopeptides, albeit with a limited scope ^44^. These platforms achieve their speed through specialized algorithms and efficient workflows, making them a valuable method of choice for the rapid screening of glycoprotein databases from an organism’s complete proteome.

This study presents a comprehensive algorithm, **G**lycoproteomics **D**ata **A**nalysis **S**oftware (**GDAS**), designed to discover disease-specific glycosylation (DSG) from MS data of the complete human proteome (**Figure 1**). The workflow begins with a rapid, broad-spectrum analysis of MS^n^ spectra, using an “open search” approach to screen raw data, identify potential glycopeptides, and reduce the extensive protein database to a more manageable subset, conserving computational resources. Subsequently, the workflow branches into specialized analyses for N-linked and O-linked glycosylation. Specific software tools are employed for each type, which use a glycan-first strategy that leverages the predictable fragmentation patterns of glycans to confidently identify and localize them on a peptide backbone. The resulting glycoprotein database is then further refined based on statistical significance using metrics such as fold-change (FC), p-value, Mean Squared Error (MSE), Root Mean Squared Error (RMSE), and R^2^. Finally, the analysis of disease-specific glycosylation markers involves integrating quantitative data from site-specific glycosylation analysis with information on disease-related signaling pathways, biological processes, and protein-protein interactions from databases like KEGG ^45^, GO ^46^, and GeneMANIA ^47^. This allows for a complete deciphering of the disease-relevant protein glycosylation.

**Figure 1.**
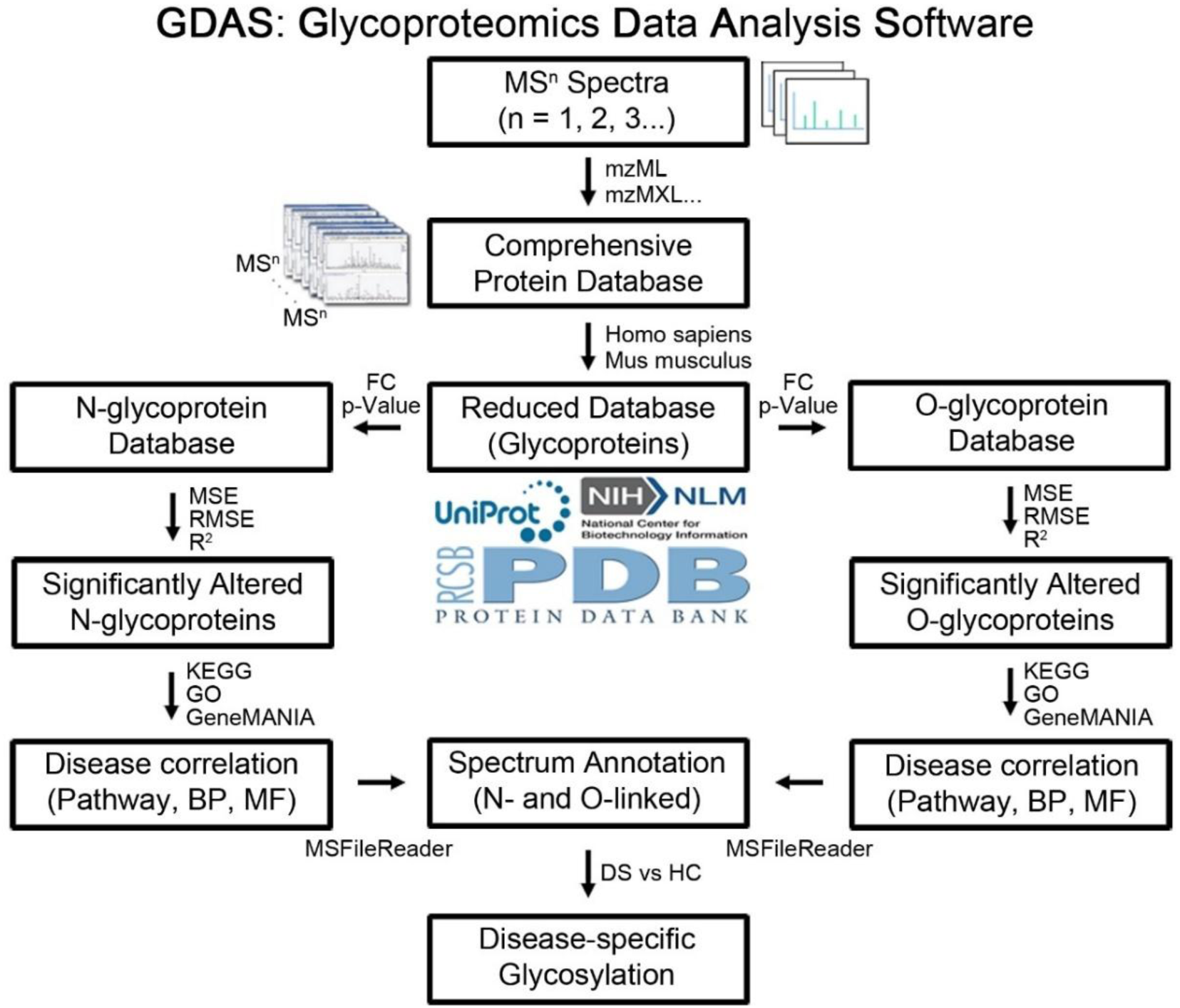
Schematic workflow of the Glycoproteomics Data Analysis Software (GDAS). This diagram illustrates the comprehensive strategy for the high-throughput identification of disease-specific protein glycosylation. The process begins with the analysis of MS^n^ spectra against a comprehensive protein database (e.g., human or mouse proteome). This initial step generates a reduced database of identified glycoproteins. This reduced database is then further filtered to include only those proteins where glycosylation is statistically significantly altered, as determined by metrics such as fold-change (FC), p-value, Mean Squared Error (MSE), Root Mean Squared Error (RMSE), and R^2^. The targeted proteins are then subjected to detailed analysis for site-specific glycosylation annotation. Finally, to pinpoint disease-specific glycosylation markers, the data is correlated with disease-related pathways, biological processes, and molecular functions using databases such as KEGG, GO, and GeneMANIA.

## Results

### GDAS schematic interface

The GDAS interface facilitates the sequential analysis of protein glycosylation (**Figure S1**); the necessary GDAS installation is detailed in the Supplementary Materials. The initial step involves using the interface to upload the protein database and MS raw data (disease cohort vs. control cohort) (**Figure S1A**), with a progress description appearing after clicking “Run”. For N-glycosylation analysis, selecting “N-glycosylation” under the “Mode” folder (**Figure S1B**) initiates MSFragger to generate a glycoprotein database for GlycReSoft analysis, which automatically uploads the MS raw data and MSFragger results (**Figure S1C**); GlycReSoft then quantitatively characterizes significantly regulated protein glycosylation and glycoforms based on selection criteria such as fold-change (FC) (e.g., >1.5) and p-Value (e.g., <0.05). Likewise, selecting “O-glycosylation” under “Mode” (**Figure S1D**) proceeds with O-glycosylation analysis by running MSFragger followed by O-Pair analysis (**Figure S1E**). Finally, the results from either GlycReSoft or O-Pair are uploaded for a final Byonic analysis, which generates detailed glycoforms and site-specific glycosylation annotation (**Figure S1F**); it’s important to note that alternative software with comparable performance may be used in place of the specified tools.

### Computation algorithm for significantly regulated protein glycosylation

The computational challenge of analyzing the entire human proteome for protein glycosylation - without prior knowledge from a targeted protein database - arises from two factors: the inherent complexity of glycosylation itself and the subsequent complicated fragmentation ions of glycopeptides observed in MS. Therefore, an efficient proteomic approach is ideally required to quickly identify proteins containing glycosylation sites without the extensive computation time needed for searching detailed site-specific glycoforms.

To achieve this goal, an efficient data analysis workflow was established, starting with an ultrafast glycoproteomic search engine followed by extensive glycoform analysis. We integrated a tool like MSFragger-Glyco ^48^, known for its fast and sensitive identification of N-linked and O-linked glycopeptides, making it suitable for the initial screening against a complete proteome database (e.g., human, mouse, dog) using raw MS data (**Figure 2** & **Figure S2**). This preliminary analysis identifies N-linked and O-linked glycoproteins, where overall glycosylation changes between disease and healthy controls are determined using FC and a p-value; these metrics are used to generate a volcano plot that reduces the protein database from over 10,000 to hundreds of statistically significant proteins. To then provide a rapid, quantitative extraction of N-glycan compositions and abundances, and to assign glycan compositions using supervised/unsupervised scoring methods from LC/MS data, GlycReSoft ^27^ is applied to further refine the protein database. Likewise, O-Pair ^28^ is utilized to define O-glycosite localization (steps 1 to 8), capable of reducing search times over 2000-fold compared to other tools.

**Figure 2.**
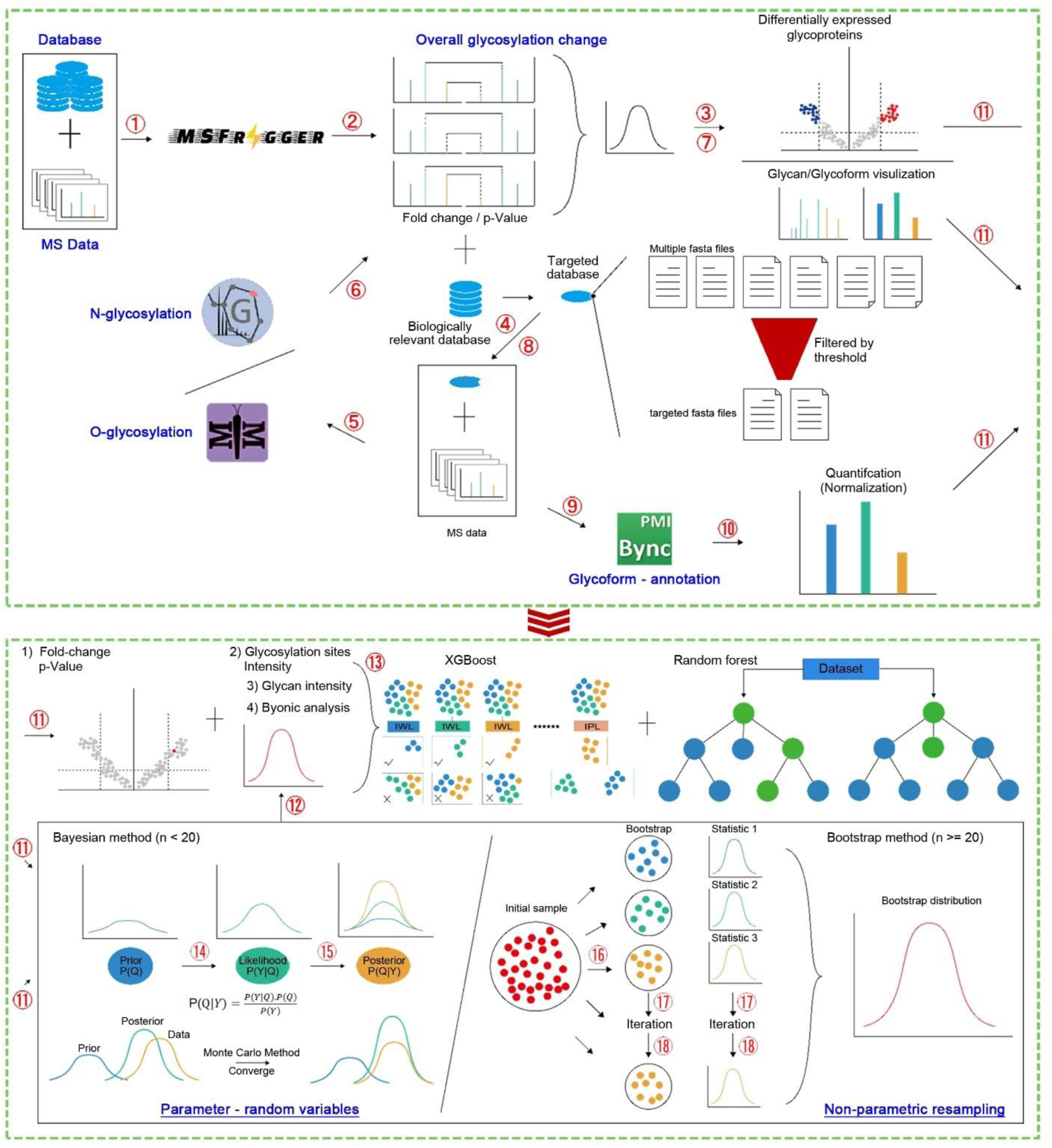
Schematic diagram of the computational algorithm for GDAS. This figure outlines the workflow for identifying protein glycosylation biomarkers, encompassing initial processing, machine learning, and statistical validation. The process begins with Initial Processing and Filtering where (**1**) MS Data and a protein database are input into MSFragger for an initial library search. (**2**) Overall glycosylation change is determined by calculating FC and p-values from these results, which are then (**3**) visualized as a volcano plot to show differentially expressed glycoproteins. A targeted database is created by (**4**) filtering the protein database based on user-defined FC and p-value thresholds. Subsequently, for a more specific analysis, (**5**) N-glycosylation and O-glycosylation analyses are performed using specialized tools (GlycReSoft for N-linked and O-Pair for O-linked) with the original MS data and the filtered protein library. (**6**) Differential glycan analysis calculates FCs for these results, and (**7**) Glycan/Glycoform visualization displays the corresponding glycosylation sites and glycan structures. This leads to (**8**) Database refinement, creating a biologically relevant, reduced protein database. In a (**9**) second pass of glycosylation analysis, this refined database and the original MS data are processed by Byonic for comprehensive (**10**) Glycoform annotation. All results, including those from the volcano plot, undergo (**11**) Data pre-processing and characterization for downstream analysis. The workflow then transitions to Machine Learning and Statistical Analysis. (**12**) Data preparation formats the pre-processed data for the XGBoost algorithm. (**13**) Biomarker identification is achieved by establishing a predictive model using XGBoost to screen for potential biomarkers. For Statistical methods for validation: (**14**) Bayesian method (n < 20) is applied when the sample size is less than 20, where a prior model is iteratively (**15**) corrected to approximate the true distribution. For sample sizes of (**16**) n ≥ 20, the Bootstrap method is employed, generating multiple sub-sample sets. (**17**)-(**18**) Distribution analysis then calculates statistics from these sub-samples, iteratively integrating and analyzing the overall dataset’s distribution.

Next, Byonic ^49^ is utilized for targeted analysis of protein glycosylation, providing detailed MS/MS fragment annotation along with other features. The fold-change (FC), p-value, intact glycopeptide intensity, and glycan abundance are statistically calculated using various statistical tools: XGBoost, Random Forest, or Bayesian methods when the sample size is less than 20, and the Bootstrap method when the sample size is 20 or greater (**Figure 2**). These statistical analyses generate a comprehensive glycosylation score based on the glycan profile, glycosylation site localization, FC, p-value, and quantification (**Figure S2**). This combined application of multi-layered glycoproteomic search tools and advanced statistical methods facilitates the precise identification of targeted glycoproteins that exhibit statistically significant disease-related changes, thereby enabling the pinpointing of potential disease-specific glycosylation markers.

### GDAS validation using model glycoprotein and complex biological sample

We used fetuin from bovine serum to compare the N-glycan and O-glycan profiles derived from the GDAS search engine with those from GlycReSoft and O-Pair analyses (**Figure 3**A). The GDAS search on intact fetuin glycopeptides identified six major N-glycans (H5N4, S1H5N4, S2H5N4, S1H6N5, S2H6N5, S3H6N5), with S3H6N5 exhibiting the highest relative abundance ^50^, a profile that was consistent with the GlycReSoft analysis. Similarly, the fetuin O-glycan profile was consistent with literature ^51^, as both the GDAS and O-Pair searches showed similar relative abundances for each O-glycan. Furthermore, we inspected the site-specific glycoforms for both N- and O-linked glycosylation (**Figure 3**B), where site-specific mapping showed highly sialylated glycans on N[99]CS and N[156]DS with 100% occupancy, but less occupancy on N[176]GS ^52^. GDAS successfully annotated MS/MS fragments with b, y, and oxonium ions, and six O-glycosylation sites were confirmed, each with a definite relative abundance and diverse O-glycoforms. These results collectively validated the accuracy of the quantitative analysis of fetuin glycosylation using the combined workflow.

**Figure 3.**
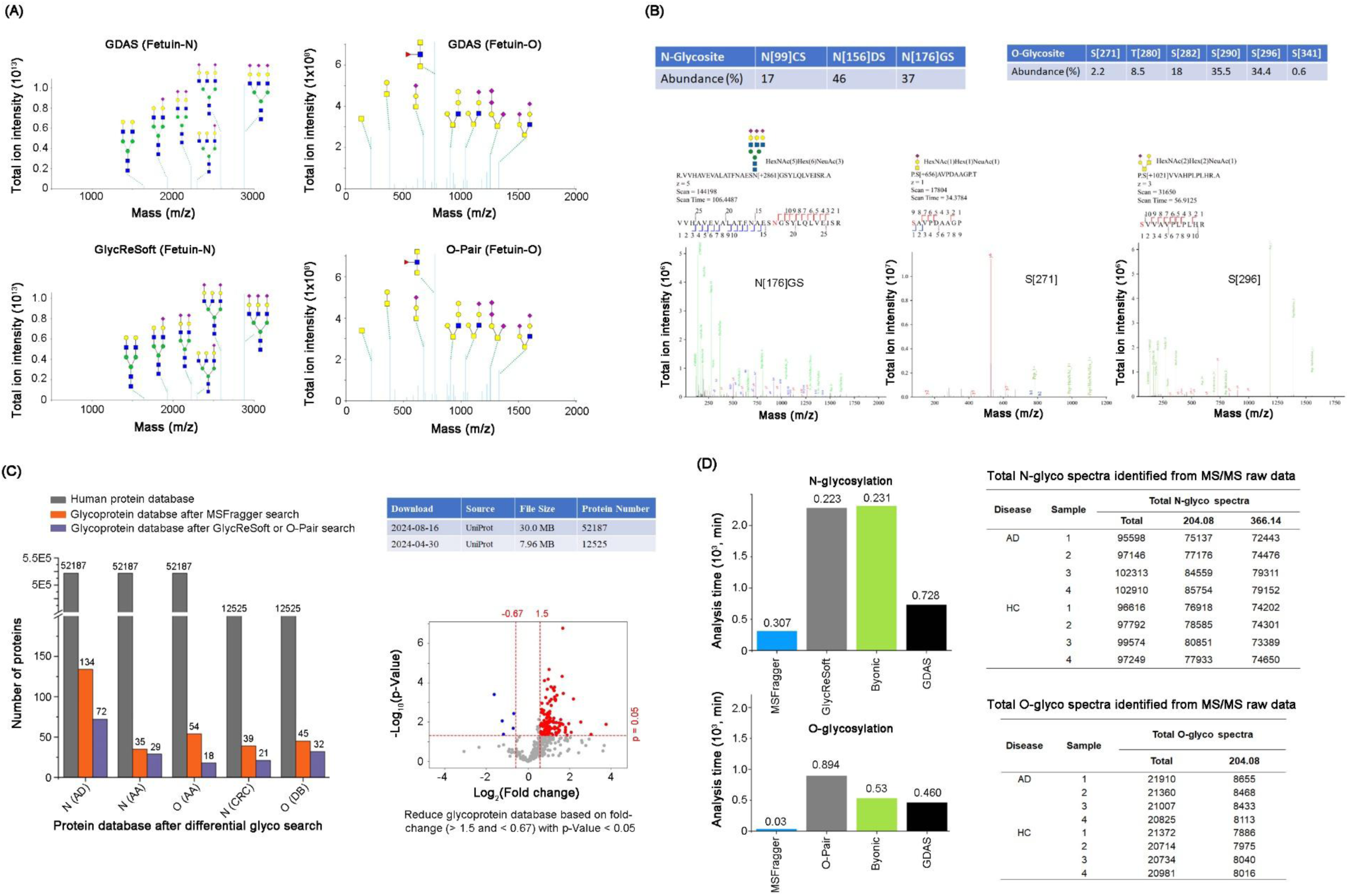
Validation and comparison of the GDAS glycosylation analysis. This figure validates the GDAS workflow using model glycoproteins and complex biological samples for both N-linked and O-linked glycosylation. (**A**) Shows the glycan profiles of bovine fetuin generated by GDAS compared to established software, GlycReSoft (for N-glycans) and O-Pair (for O-glycans). The similar type and relative abundance of glycans across these analyses demonstrate the accuracy of the GDAS approach. (**B**) Provides a detailed MS/MS annotation of fetuin glycosylation sites and their attached glycans, specifically characterizing the relative abundance of three N-glycosites and six O-glycosites. Representative spectra for the N[176]GS, S[271], and S[296] sites are shown to illustrate the confident identification. (**C**) Demonstrates the step-wise reduction of the protein database in GDAS analysis for various diseases. Starting with a large human proteome database (52,187 proteins), the analysis on Alzheimer’s Disease (AD) samples shows the database size is reduced to 134 proteins after an MSFragger search and further narrowed to 72 proteins with significantly altered glycosylation. Similarly, for asthma (AA), the database is reduced from 52,187 to 35 proteins for N-glycosylation and 54 for O-glycosylation after MSFragger, and further refined to 29 and 18 proteins, respectively, after specific glycosylation searches. Using a smaller human proteome database (12,525 proteins), the number of proteins is reduced for N- and O-linked analysis for colorectal cancer (CRC) and diabetes, respectively. This database reduction is driven by statistical analysis, as visualized in the volcano plot, which highlights significantly altered glycoproteins. (**D**) Compares the analysis time of different software tools with the GDAS workflow. MSFragger is shown to be the fastest for initial N- and O-glycosylation analysis using MS data from four AD and four healthy control samples. The GDAS workflow integrates the strengths of these individual software tools to optimize both speed and comprehensive analysis.

We used data from our previous published work and papers from other laboratories to demonstrate the effective reduction of the protein database achieved by the GDAS workflow. For example, when using the MS spectra of AD glycopeptides ^53^, GDAS essentially reduced the human protein database from 52,187 to 134 after an MSFragger search, which was further reduced to 72 by either GlycReSoft or O-Pair searches (**Figure 3**C). Similarly, MS spectra of N-glycopeptides generated from the sera of patients with asthma versus healthy controls (HC), when searched against the human proteome, were effectively reduced from 52,187 to 35 by MSFragger, and further to 29 by GlycReSoft (N-glycosylation). In parallel, MS spectra of O-glycopeptides derived using the MOTAI method from the same patient group showed a significant reduction of the protein database when considering FC and p-value. Furthermore, tissue specimens from colorectal cancer (CRC) patients ^54^ compared to paracancerous controls contained 39 significantly different glycoproteins, with 21 targeted N-glycoproteins for Byonic analysis; here, GDAS similarly reduced the number of serum proteins of Diabetes patients (DB)^55^ from 12,525 to 45 by MSFragger and further to 32 by O-Pair, highlighting the workflow’s effectiveness in identifying a targeted protein database by utilizing FC and p-value, as depicted in the volcano plot.

The GDAS workflow can significantly reduce the analysis time required for glycoproteomics studies. For instance, in an N-glycosylation analysis using four MS spectra from AD and four from HC - with typical oxonium ions at m/z 204.08 and 366.14 - searching the entire human protein database (52,187 proteins) against a human glycome database using MSFragger took 307 minutes (**Figure 3**D). In comparison, searching the same entire protein database for these MS spectra took approximately 2,278 minutes by GlycReSoft and 2,310 minutes by Byonic. Crucially, when the GDAS algorithm was employed, it took only 728 minutes to identify the targeted glycoproteins with site-specific glycoforms, demonstrating a substantial time saving. Furthermore, since current commercial software cannot directly identify significantly regulated protein glycosylation, especially O-glycosylation, by searching against the entire human proteome, GDAS was specifically developed to address this critical challenge.

### Identification of disease-specific glycosylation markers in complex biological sample

The MS spectra data were downloaded from the ProteomeXchange repository (ID: PXD022274), which included four AD and four HC tissue samples. Protein database (52,187 proteins) from Uniprot and human glycan databases from CFG, Carbbank, GlycomeDB, and Glycosciences (**Table S1**) were also obtained. The MS spectra were initially processed using MSFragger to identify N- and O-linked glycosylated proteins and calculate the FC and p-value between AD and HC, visualized in a volcano plot (**Figure 4**A). Proteins meeting the criteria of FC > 1.5 or < 0.67 and p < 0.05 were selected to create a reduced protein database for subsequent analysis by GlycReSoft, significantly decreasing computation time. GlycReSoft then used the N-glycan database on these selected proteins to generate a plot of FC for each identified glycoprotein. This FC information was, in turn, used to further refine the protein database for a Byonic search, where metrics like glycan intensity, intact glycopeptide intensity, and glycosylation site were measured to calculate R^2^, MSE, and RMSE, ultimately yielding a glycosylation score for each protein. This final score was used to rank the potential importance of each protein in AD, effectively filtering out proteins with potentially less impact on the disease pathology.

**Figure 4.**
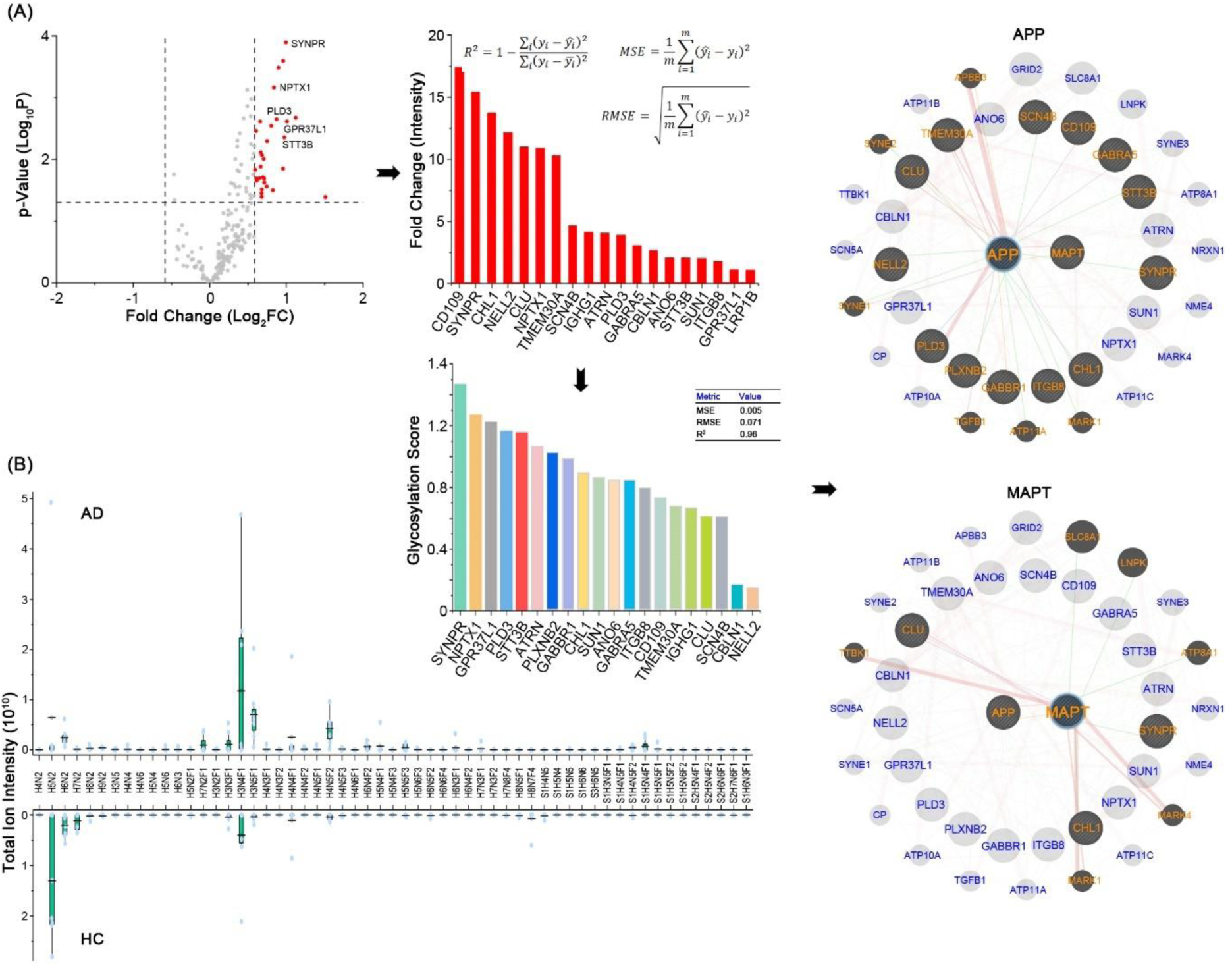
Identification of disease-specific N-glycosylation from AD tissues using GDAS. This figure presents the results of the GDAS analysis on Alzheimer’s disease (AD) tissue samples. (**A**) The analysis begins by processing MS data from four AD and four healthy control (HC) samples against a human proteome database and known human glycome. The volcano plot displays the distribution of glycoproteins based on p-value and fold-change (FC), with statistically significant hits highlighted in red. The top-ranked glycoproteins by FC are shown, with their significance confirmed by R^2^, MSE, and RMSE values. The proteins with the highest glycosylation scores are then identified through Byonic analysis. To understand their biological relevance, these proteins are analyzed using databases such as KEGG, GO, and GeneMANIA and are mapped onto key AD signaling pathways, including amyloid-β and tau. The figure highlights the protein-protein interaction networks of two central AD proteins, amyloid-β precursor protein (APP) and microtubule-associated protein tau (MAPT), with the significantly altered glycoproteins. (**B**) Compares the overall N-glycome profile between AD and HC samples, showing the total ion intensity of various glycan compositions. The nomenclature used is H (Hexose), N (HexNAc), F (Fucose), S (Neu5Ac), and G (Neu5Gc).

Based on the calculated glycosylation score, the glycoproteins ranked by importance, from most to least, are SYNPR, NPTX1, GPR37L1, and PLD3, among others. To more thoroughly assess which of these glycoproteins potentially impact AD pathogenesis, their involvement in known signaling pathways and molecular processes must be considered. For example, the amyloid cascade hypothesis proposes that the cleavage of amyloid precursor protein (APP) results in amyloid aggregation, a key mechanism in AD ^56^, while more recent studies on amyloid-β (Aβ) signaling have highlighted differential interactions between Aβ and Tau-mediated changes, neuroinflammation, or other inflammatory processes ^57^. Therefore, these observed abnormal alterations in protein glycosylation should be correlated with an imbalance in Aβ signaling. To elaborate on this correlation, the identified glycoproteins were mapped using GeneMANIA ^47^, to establish their associations with APP and MAPT (microtubule-associated protein Tau), as illustrated in **Figure 4**. The resulting protein-protein interactions show that APP correlates with several AD-related glycoproteins, including CLU, CHL1, and SYNPR, and similarly, MAPT also interacts with these glycoproteins, strongly suggesting that CLU, CHL1, and SYNPR have significant implications in AD pathogenesis.

The GDAS also generated the comparative glycan profiles between the Alzheimer’s disease (AD) and healthy control (HC) groups, which are presented in **Figure 4**B. These profiles were categorized by glycan type, including high-mannose, hybrid glycans, fucosylated complex glycans, sialoglycans, and fucosyl-sialoglycans. Given the use of multiple samples, the standard deviation was calculated and indicated within the profile. This visualization demonstrates the distinct changes in N-glycans observed in AD, thereby providing critical insight into which glycoenzymes may be altered in the disease.

### AD-specific glycosylation marker identification from CSF

For the purpose of early diagnosis, tissue-specific markers are generally less ideal than those found in body fluids, which is why we utilized MS spectra from the CSF of patients with AD versus HC to investigate elevated protein glycosylation and determine if CSF changes correlate with tissue alterations. The GDAS readily identified elevated N-glycosylation to generate a glycosylation score, which highlighted CSF-specific markers such as MRCAM and CADM2 (**Figure 5**A). MAPT shows a correlation with both CADM2 and MRCAM, noting that the *CADM2* gene may influence neurocognitive functions ^58^ and MRCAM serves as a marker for the substrate-selective activation of ADAM10 in AD ^59^. Furthermore, the N-glycan profile analysis of the CSF revealed a noticeable decrease in core-fucosylation and an increase in high-mannose glycans in the AD group (**Figure 5**B).

**Figure 5.**
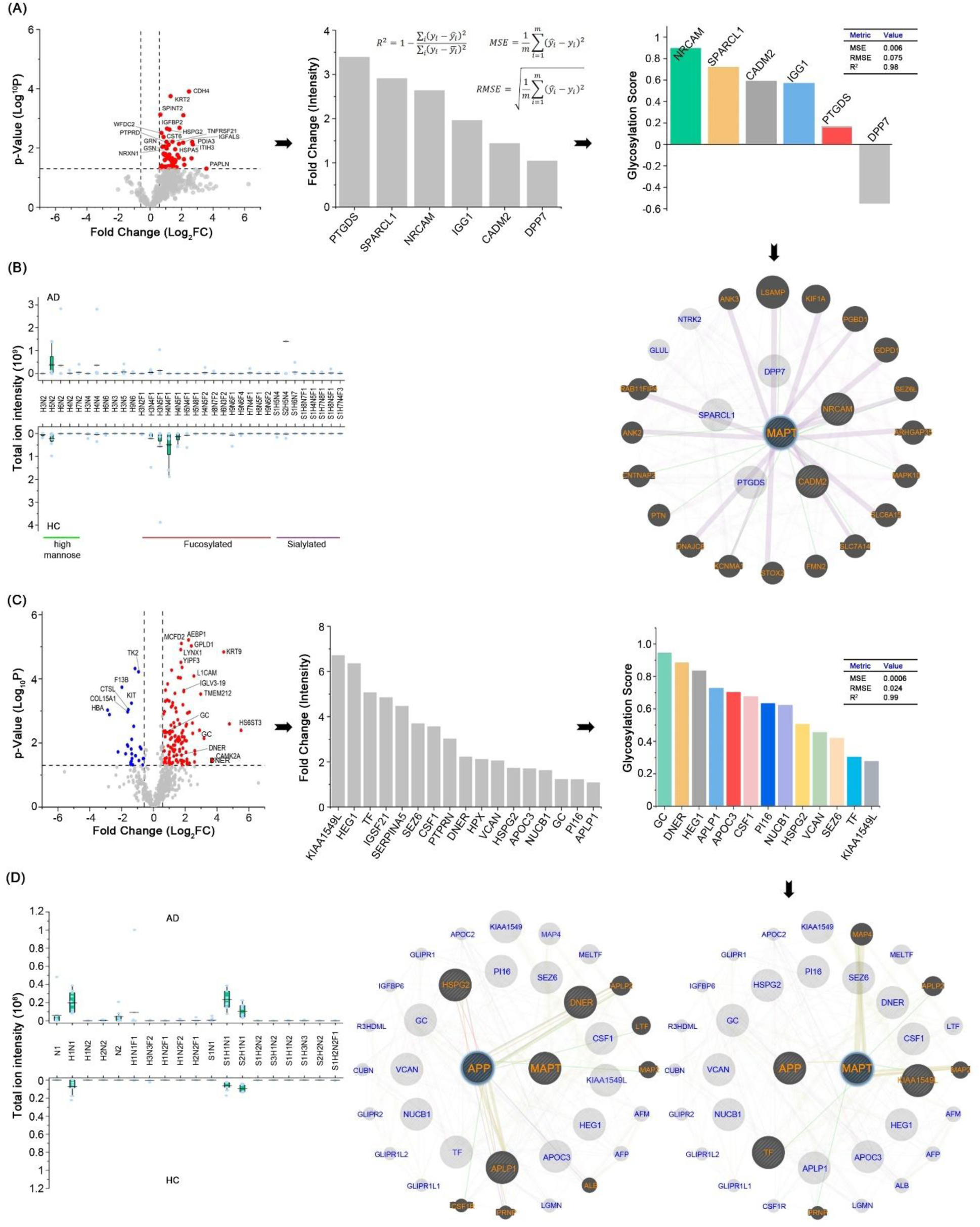
Identification of potential cerebrospinal fluid (CSF) protein N-linked and O-linked glycosylation markers using GDAS. This figure presents the analysis of both N-linked and O-linked glycoproteins in cerebrospinal fluid (CSF) samples from Alzheimer’s Disease (AD) patients compared to healthy controls (HC). (**A**) The volcano plot generated by MSFragger-Glyco highlights significantly upregulated N-glycoproteins in AD, including PTGDS, SPARCL1, NRCAM, IGG1, CADM2, and DPP7. These proteins are further analyzed by GlycReSoft and Byonic to determine their glycosylation score. Notably, NRCAM and SPARCL1 show high glycosylation scores, indicating their significant alteration in AD. A protein-protein interaction network shows that NRCAM and CADM2 are associated with MAPT (Microtubule-Associated Protein Tau), suggesting a link to AD pathogenesis. (**B**) Compares the overall N-glycome profile between AD and HC, revealing an upregulation of high-mannose structures and a downregulation of fucosylated glycans in AD samples. (**C**) A separate volcano plot identifies significantly upregulated O-glycoproteins, such as KIAA1549L, HeG1, TF, and IGSF21. GC, NDER, HEG1, and APLP1 exhibit particularly high glycosylation scores. Further pathway analysis reveals that HSPG2, APLP1, and DNER interact with APP (Amyloid-β Precursor Protein), while TF and KIAA1549L are correlated with MAPT. (**D**) Compares the O-glycome between AD and HC, showing a specific upregulation of O-glycosylation on T antigen (H1N1), sialyl T antigen (S1H1N1), and S2H1N1 in AD.

The GDAS analysis focusing on O-glycosylation in CSF identified a substantial increase in O-glycoproteins (with a FC of > 1.5 and a p-value of < 0.05). An O-Pair search was performed, revealing the FC for the top 19 O-glycoproteins, with the glycosylation score ranking GC (group-specific component), DNER (delta and notch-like epidermal growth factor-related receptor), and HEG1 (HEG homolog 1) among the highest (**Figure 5**C). Considering their known roles with APP and MAPT (Tau), the glycoproteins HSPG2 (heparan sulfate proteoglycan 2), DNER, and APLP1 may have a particularly high impact on AD pathogenesis, while TF and KIAA1459L also show interactions with MAPT. The comparative O-glycan profile derived from these glycoproteins, as shown in **Figure 5**C, notably revealed an increase in both T antigen and sT antigen.

### Annotation of MS/MS fragment

We selected several key glycoproteins for illustration based on the GDAS results and their documented co-presence in both tissue and CSF, as summarized in the Venn diagram of (**Figure 6**), which highlights those exhibiting increased expression in AD. Focusing on their relationship with APP and MAPT, the AD-associated glycoproteins CADM2, NRCAM, CLU, and CHL1 were extensively annotated via their MS2 spectra (**Figure 6**B and **6**C), where the structure of the intact glycopeptide is determined using characteristic B-ions (oxonium ions) and Y-ions. The GDAS output provides comprehensive molecular details, including the glycan composition, intensity, peptide sequence, charge, scan time, and a cartoon visualization of the intact glycopeptide. For instance, CADM2 and MRCAM from CSF show varied glycoforms between AD and HC samples, while the tissue-specific glycoprotein CLU exhibits higher levels of Man5 (H5N2) and biantennary sialoglycan (S2H5N4) in AD compared to higher expression of Man9 (H9N2) and S1H5N5F1 in HC; similarly, CHL1 presents different glycoforms in AD versus HC tissue, as studies have shown that CHL1 concentration in the CSF correlates with Tau and phosphorylated Tau ^60^. These findings confirms that GDAS enables the precise, site-specific identification of these altered glycoforms in AD.

**Figure 6.**
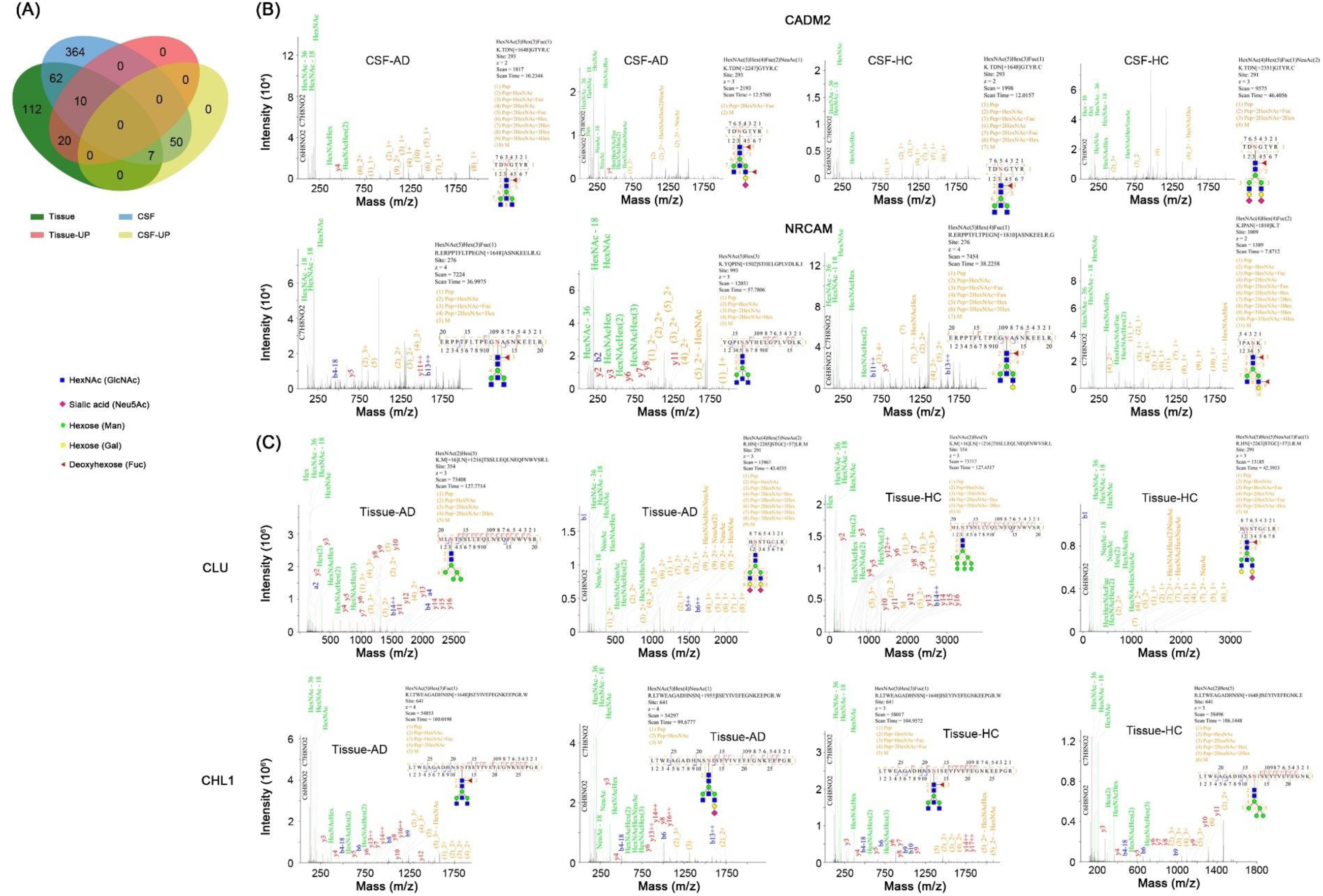
Identification of site-specific glycoforms using annotated tandem mass spectrometric fragment ions. (**A**) This Venn diagram compares the identification of glycoproteins found in tissue and CSF, highlighting the subset of those glycoproteins that are upregulated in AD (labeled as Tissue-UP and CSF-UP). (**B**) The figure presents the tandem mass spectrometric (MS2) annotation of intact glycopeptides for CADM2 and NRCAM in CSF, comparing AD and Healthy Controls (HC). The annotated fragment ions, including peptide + oxonium ions, provide detailed information distinguishing site-specific glycoforms, indicating differences in intact glycopeptide structures between conditions. (**C**) Similarly, this panel shows the tandem mass spectrometric annotation of intact glycopeptides for CLU and CHL1 in Tissue, comparing AD and HC samples.

## Discussion

High-throughput analysis of complex N- and O-glycosylation faces significant computational challenges, primarily due to the vast heterogeneity of glycan structures and the need to process mass spectrometry data against large protein and glycan databases. While UniProtKB/Swiss-Prot contains approximately 20,000 reviewed unique protein-coding genes and over 22,000 additional isoforms, hundreds of glycans have been identified from human cell lines, body fluids, and tissues ^61,62^. Furthermore, different MS/MS fragmentation techniques produce a variety of ions, such as a, b, c, x, y, z, or B-ions/Y-ions (**Table S2**), complicating the matching process. Calculating the masses of these diverse fragment ions against both the large protein and complex glycan databases results in extremely lengthy computation times, making it currently impractical to derive disease-specific glycosylation markers with detailed glycoforms. Therefore, a new computational strategy is required to resolve this analytical challenge.

Here we developed an algorithm, termed GDAS, to expedite the analysis of protein glycosylation using MS spectra alongside large protein and glycan databases. GDAS initiates the process by integrating two groups of multiple MS spectra data, the human proteome database, and a database of known N- and O-linked glycans (**Figure S2**). The algorithm first employs a rapid analysis to identify proteins that are glycosylated, substantially reducing the subsequent search space for the next module. Following this, statistical analysis is performed to generate FC and p-values for each identified glycoprotein, leading to the selection of statistically significantly regulated protein glycosylation events. This initial filtering strategy drastically decreases the protein database size when interfaced with downstream software for quantitative analysis. Specifically, if N-glycosylation is considered, any compatible software providing detailed quantitative analysis of the intact glycopeptide can be used (**Table 1**); similarly, software capable of analyzing O-glycosylation can then be utilized to further target specific O-glycoproteins.

The GDAS software utilizes a multi-step statistical and machine learning pipeline in its final analysis module, beginning with statistical evaluation of four key data matrices (FC/p-value, glycan intensity, site-intensity, and Byonic results) using either the Bayesian method for smaller sample sizes or the Bootstrap method for larger sample sizes (**Figure S5**, **S6** and **S8**). The Bootstrap method generates numerous surrogate datasets by resampling with replacement to accurately examine data variability and uncertainty, while the Bayesian method updates probability estimates using a robust Cauchy prior distribution *via* the Markov Chain Monte Carlo (MCMC) method to calculate glycan and glycosylation sites distribution. The statistical outputs from these two methods are then used as eigenvalues for two machine learning algorithms: XGBoost (eXtreme Gradient Boosting) and Random Forest. XGBoost, an optimized Gradient Boosting Decision Tree (GBDT) framework, is employed to build a non-linear regression model that uses the fold-change matrix as the dependent variable to evaluate the effect of glycosylation on expression, resulting in a weighted score matrix. In parallel, the Random Forest ensemble algorithm is used for classification and feature correction, leveraging the bagging concept to adjust the values of each statistic and form the final statistic matrix for subsequent Principal Component Analysis (PCA).

Considering the interplay of biological processes and molecular functions, including disease-involved signaling pathways, it is crucial to screen for the most relevant disease markers. This requires establishing the disease-involved pathways by analyzing vast biological datasets, such as genomic, proteomic, and transcriptomic data, to identify the patterns and interactions that reveal the underlying molecular mechanisms and dysregulated pathways. Through data integration, it is ideal to generate protein-protein networks and utilize a systems biology approach, performing pathway enrichment analysis using databases like GO, KEGG, or STRING. Correlating these abnormal biological processes (e.g., glycosylations) with the dysregulated pathways can significantly aid in the identification of disease-specific markers. In our ongoing work, we will integrate these components into the GDAS.

## Conclusion

The Glycoproteomics Data Analysis Software (GDAS) represents a significant advancement in glycoproteomics by offering a powerful, high-throughput platform for the comprehensive identification of disease-associated glycoforms. A primary advantage of GDAS is its ability to overcome the major computational challenges associated with searching an entire proteome database for protein glycosylation, which is typically time-consuming and demanding. The workflow’s core technical finding is an efficient, multi-step strategy: it begins with a rapid, broad-spectrum “open search” to screen for significantly regulated glycosylation, which drastically reduces the extensive protein database to a manageable subset. This data reduction was validated across multiple complex biological samples, where the massive human protein database of over 52,000 entries was successfully filtered down to a few dozen statistically significant targets in studies of Alzheimer’s disease, asthma, and diabetes. The platform’s overall accuracy and quantitative analysis capabilities were also confirmed by demonstrating that its N-glycan and O-glycan profiles for the model glycoprotein fetuin were consistent with established software and literature.

The usefulness of GDAS is amplified by its sophisticated quantitative analysis and integration of multiple specialized search tools for both N-linked and O-linked glycosylation. The Final Analysis Module is particularly useful, as it utilizes a robust statistical and machine learning pipeline, incorporating methods like Bootstrap/Bayesian statistics and algorithms such as XGBoost and Random Forest. This advanced computational approach corrects for factors, mitigates multicollinearity, and generates a comprehensive, weighted glycosylation score. By combining this efficient data reduction strategy with powerful multi-layered analysis, GDAS facilitates the precise identification of statistically significant, disease-specific glycoforms. The final utility of the platform is realized through its ability to integrate quantitative glycosylation data with information from external databases (e.g., KEGG, GO). This allows for the complete deciphering of disease-relevant protein glycosylation, making GDAS an invaluable, high-throughput tool for accelerating biomarker discovery.

## Supporting information

Installation guide

## Online content

All methods, additional references, summaries, extended data, supplementary information, and statements of data and code availability, including details of author contributions and competing interests, are accessible online.

## Data availability

The LC-MS/MS data have been deposited to the ProteomeXchange Consortium via the MassIVE partner repository with the dataset identifier PXD022274.

## Ethics Approval

Approval for this study was obtained from the ethics committee of the Second Affiliated Hospital of Soochow University (JD-LK2023001-R01). Written informed consent was secured from all participating patients.

## Acknowledgments

We thank the Jiangsu Science and Technology Plan Funding (BX2022023), the Jiangsu Shuangchuang Boshi Funding (JSSCBS20210697), the Suzhou Medical Innovation Funding (SKJY2021141), and the Frontier Discipline - Pediatric Respiratory Medicine (ML13101123) for their generous support.

## Author contributions

W.S.Y., G.Y., M.X.Y., D.J.B., and Z.Y.F. conducted the experiments and data analysis, prepared the standard operation procedure, and compiled the video. G.W. performed the GO and KEGG analysis. B.S.Y., Z.W.Q., and Z.R.M. were responsible for preparing samples for MS analysis.

H.C.Y. and M.J.F. generated the LC-MS/MS data. J.H.J. provided funding support. S.Y. designed the study, drafted the manuscript, and also provided funding support. All authors edited and reviewed the manuscript.

## Supporting information

Supporting information includes detailed sections on the Total Analysis Module, Final Analysis Module, statistics methods, machine learning algorithm, and prediction logic, complemented by the following tables and figures: Table S1 (Glycan database), Table S2 (Fragmentation ions), Figure S1 (GDAS interface), Figure S2 (GDAS algorithm), Figure S3 (GDAS procedure), Figure S4 (Machine learning), Figure S5 (Bootstrap), Figure S6 (Bayesian), Figure S7 (GDAS diagram), Figure S8 (XGBoost), and Figure S9 (MS2 annotation).

## Methods

### GDAS search algorithm

The GDAS workflow employs a two-pronged strategy to efficiently analyze glycoproteomics data. Initially, in the Total Analysis Mode, GDAS selects the appropriate module (N- or O-glycosylation) and pre-processes the MS spectra based on defined quantitative thresholds (e.g., FC > 1.5 and p-value < 0.05). Concurrently, the fold change (FC) and p-value for proteins between the two analyzed groups are calculated. The pre-processed MS spectra are imported into the integrated MSFragger tool for classification pre-processing (**Figure 2** and **Figure S2**). Simultaneously, a comprehensive protein database is downloaded from UniProt (e.g., Homo sapiens or Mus musculus) ^63^. To counter the inherent False Discovery Rate (FDR) in p-value calculations, the system strictly selects only proteins that satisfy the user-defined FC and p-value thresholds for statistical significance. Proteins that are identified by MSFragger as having high FC values and verified N-linked and/or O-linked glycosylation are then exported to create a highly focused, targeted glycoprotein database. This initial step is critical, as it significantly reduces the protein search space, enabling a much faster and more efficient glycosylation analysis under the “Total Analysis Mode” compared to searching the entire comprehensive protein database.

The glycoprotein database generated in the initial step of GDAS is integrated with more specialized glycosylation analysis software, such as GlycReSoft for N-glycosylation or O-Pair for O-glycosylation. By selecting the appropriate analysis with “**N/O-Glycosylation Analysis Mode”**, GDAS leverages the same MS spectra to comparatively analyze the differential expression of N-linked and O-linked glycoproteins, respectively. User-defined criteria, such as the fold-change of glycoproteins between disease and healthy control groups, are applied during this step to further refine the target glycoprotein list. The total ion abundance of glycans and glycoforms associated with each glycoprotein can be visualized and exported as an Excel file, enabling the identification of dysregulated glycoproteins for subsequent site-specific glycosylation analysis.

Following the “**N/O-Glycosylation Analysis Mode**”, targeted glycoproteins are analyzed using Byonic software. To assess the biological significance of changes in disease-specific glycosylation, statistical parameters can be incorporated to demonstrate significance. Byonic offers various search strategies, including glycan-first, peptide-first, or combined glycan-peptide approaches, to efficiently, accurately, and precisely identify intact glycopeptides. This powerful tool provides detailed annotations for fragment ions, encompassing peptide backbone ions (a, b, c, x, y, z, and Y-ions) and glycan-related ions (B-ions, oxonium and glycan fragment ions). However, using a comprehensive protein database for disease-specific glycosylation analysis is impractical due to the excessive time required to process mass spectra data. To address this, GDAS pre-identifies glycoproteins of interest, enabling a more focused and efficient Byonic analysis using a limited set of glycoproteins. This strategy significantly reduces the search time and minimizes the identification of non-glycoproteins or subtle glycoprotein changes, ultimately improving the overall efficiency of the analysis. Upon completion of the search, Byonic outputs site-specific glycosylation data.

The core of the “**Final Analysis Mode**” prediction algorithm involves utilizing Bootstrap to quantify glycoforms (**Supporting Information**), glycosylation sites, glycoprotein expression matrices, and the final Byonic result matrix for each glycoprotein. These data are consolidated into four-dimensional vectors, which are then subjected to linear regression using the Ridge Regression machine learning algorithm (**Supporting Information**). The Bootstrap method aims to approximate the statistical model of the sample through iterative resampling. By repeatedly selecting a subset of samples with replacement, the law of large numbers is emulated, effectively simulating an infinite number of samples. After numerous iterations, the resulting statistical data, expressed as confidence intervals, are analyzed to provide a robust and reliable prediction.

### Calculation of analysis time

The computation time experiment is designed to compare the analysis time, defined as the cumulative time required to process all independent MS data files. This calculation is first established by determining the baseline time for each search software—MSFragger, GlycReSoft, and Byonic—when run independently. For this baseline measurement, each software starts with the same, full protein database and searches against the identical set of raw MS data files. The total time for each software (T_MSFragger_, T_GlycReSoft_, and T_Byonic_) is the sum of the time consumed during its sequential retrieval/search against the MS data files within the database. The second part of the procedure measures the time for the GDAS, which involves running the three software programs sequentially to leverage the results of previous steps. The total GDAS analysis time (T_GDAS_) is the sum of the time consumed in its three ordered steps (T_1_ + T_2_ + T_3_). Step 1 uses MSFragger on the original protein database, producing a time T_1_ and a reduced database 1. Step 2 then runs GlycReSoft against the MS data, but is constrained to the reduced database 1, yielding time T_2_ and generating a further reduced database 2. Finally, Step 3 runs Byonic against the MS data, restricted to reduced database 2, resulting in time T_3_. The crucial difference from the baseline calculation is this progressive reduction of the search space, which aims to reflect the efficiency of the tandem analysis.

### GDAS search methods

GDAS leverages a modular workflow for glycoproteomic analysis, employing MSFragger, GlycReSoft, and Byonic for the identification of N-glycosylation biomarkers, and a combination of MSFragger, O-Pair, and Byonic for O-glycosylation biomarker discovery (**Figure 1**). The initial MS spectra data, which can be generated by various instruments using fragmentation methods like HCD, CID, EThcD, or EAD, must first be converted from their proprietary RAW format to the standard mzML format. This conversion is executed using the MSConvert tool (specifically version 3.0.25329 x86_64-bit) from ProteoWizard ^64^, utilizing its centroiding and default parameters. The specific analytical parameters required for MSFragger, GlycReSoft, O-Pair, and Byonic are detailed within the documentation of each respective software.

### MSFragger search method

The glycoproteomic search strategy in MSFragger uses a high-resolution approach by setting the Precursor Mass Tolerance to 10 ppm (with Isotope Error enabled) and the Fragment Mass Tolerance to 20 ppm or 0.02 Da. The search is typically set to Fully Tryptic cleavage with up to 3 missed cleavages to account for cleavage hindrance near the bulky glycan. Fixed modifications include carbamidomethyl (+57.021464 Da) on C, and variable modifications include oxidation on M (+15.994915 Da), acetyl (Protein N-term), and crucially, deamidation on N/Q (+0.984016 Da) for N-glycan identification. Glycan assignment is performed in a subsequent targeted search by specialized modules. The initial MSFragger search focuses only on identifying the peptide backbone. Following this, PTM-Shepherd uses a library containing 182 N-glycans to assign N-glycans to Asparagine (N) sites (where N is any amino acid except Proline), while O-Pair uses a library containing 78 O-glycans to assign O-glycans to Serine (S) or Threonine (T) sites. Final results are validated using the Target-Decoy Approach to ensure both Peptide and Protein False Discovery Rates (FDR) are < 1%.

### GlycReSoft search method

The GlycReSoft search was initialized using a FASTA file generated by GDAS from the MSFragger analysis of both experimental and control groups, establishing a common protein database for subsequent glycopeptide identification. This search was performed using GlycReSoft (v0.4.24) via a Docker (v4.40.0) command line or manually through the GUI (v0.4.25) ^65^. The hypothetic glycans were strictly defined either by rules in a text file located in the cache folder or, when using the GUI, by default limits of each monosaccharide (e.g., Hex 3–9, HexNAc 2–8, Fuc 0–4, NeuAc 0–4), opening the identification space for N-glycopeptides. Since different experimental MS data required distinct search parameters, tissue samples from Alzheimer’s disease (AD) were searched using HCD fragmentation with a 20-ppm mass tolerance for both precursor and fragment ions, while cerebrospinal fluid (CSF) samples from AD, with N-glycosylation, were searched using EThcD with a 10-ppm precursor tolerance and 0.01 Da fragment tolerance. Both sets of searches included carbamidomethyl (C) as a fixed modification (+57.021464 Da) and oxidation (M) (+15.994915 Da) and deamidation (N/Q) (+0.984016 Da) as variable modifications, with the maximum number of variable modifications per peptide set to the default of three.

### O-Pair search method

The O-Pair targeted search utilized a protein database derived from the shared results of MSFragger analysis across both experimental and control groups. The predefined glycan library (in text format) was input to the O-Pair module ^28^, which performs searches separate from MSFragger or GlycReSoft due to differences in their algorithms and logic. The overall search employed a dual-fragmentation strategy: HCD was used with a mass tolerance of 20 ppm for both precursor and fragment ions, while EThcD searches were set to a tighter 10 ppm precursor tolerance and 0.01 Da fragment tolerance. Fixed modifications included Carbamidomethyl (C) (+57.021464 Da), and variable modifications included Oxidation (M) (+15.994915) and Deamidation (N, Q) (+0.984016 Da), with the maximum number of modifications per peptide set to 2. All O-Pair tasks were deployed and executed using MetaMorpheus within a Docker (v4.40.0) environment or manually through the GUI (v1.0.5).

### Byonic search method

The final Byonic search utilized a protein database established from the FASTA file regenerated by GDAS, which consolidated common identifications from both experimental and control groups after GlycReSoft/O-Pair analysis. To ensure consistency, mass tolerances and modifications were set to match the previous search parameters. For analyses involving AD tissue and CSF, a total of two common modifications and one rare modification were permitted per identified peptide, while the O-glycosylation search on CSF specifically allowed one common modification and one rare modification. The search output was rigorously filtered to achieve a peptide-spectrum match (PSM) FDR of < 1% using the FDR 2D score, which is a Byonic-computed value based on a two-dimensional target-decoy strategy that simultaneously estimates and controls both PSM and protein-level False Discovery Rates.

### GDAS annotation of tandem MS fragmentations

The GDAS program provides key identification data, primarily the Scan Number within the MS file, for peptides containing fucose, sialic acid, or high-mannose modifications. To generate an initial visualization, the built-in MSFileReader library (Thermo Fisher) uses this Scan Number to retrieve the corresponding retention time and spectral data from the raw MS file, automatically plotting the spectra and displaying basic information on the associated glycans and peptides (**Figure S9**). However, for complete and publication-ready spectral annotation, manual refinement is necessary: the MSFileReader library is again used to retrieve the required data via the Scan Number, but the image is then re-plotted in Adobe Illustrator. Within Illustrator, each relevant peak is manually identified and labeled, and the specific glycan information derived from the GDAS output is strategically incorporated to produce the final, fully annotated spectra image.

