## Supplemental methods-Figures and Tables for "A Novel Glycoproteomics Platform for High-Throughput Identification of Disease-Associated Glycoforms"

### Table of Contents

**Supporting Methods** ... 4

**Total Analysis Module for MS spectra analysis** ... 4

**The Total Analysis Module outputs data for the N/O-Glycosylation Module.** ... 4

**Final Analysis Module with XGBoost and Random Forest algorithms** ... 5

**Statistics methods** ... 6

**Independent two-sample t-test** ... 6

**Bootstrap method for sample size under 20** ... 7

**Bayesian method for sample size over 20** ... 8

**Composition of input of overall** ... 8

**Machine Learning Algorithm** ... 9

**Linear regression and XGBoost** ... 9

**Classification and regression using Random Forest.** ... 10

**The logic of prediction** ... 11

**Feature extraction and engineering** ... 11

**Statistical analysis and predictive modelling of glycosylation** ... 12

**Mass spectrometric analysis of intact glycopeptides** ... 13

**References** ... 14

**Supporting Tables and Figures** ... 17

**Table S1. The glycan databased used in intact glycopeptide analysis by GDAS.** ... 17

**Table S2. Typical fragmentation ions produced by different MS/MS techniques.** ... 33

**Table S3. Nanoflow liquid chromatography gradient.** ... 35

**Table S4. The oxonium ions used in tandem mass spectrometry analysis.** ... 36

**Figure S1. The interface and workflow of GDAS for high-throughput identification of disease-specific glycosylation.** ... 37

**Figure S2. The streamlined algorithm of GDAS analysis of protein glycosylation.** ... 38

**Figure S3. GDAS algorithm procedure.** ... 39

**Figure S4. GDAS machine learning procedure.** ... 42

**Figure S5. Bootstrap resampling and effect size quantification.** ... 44

### Supporting Methods

**Total Analysis Module for MS spectra analysis**.

The Total Analysis Module is designed for obtaining all glycoproteins from MS spectra in complex biological samples. It consists of two functional sub-modules within GDAS: one for extracting common proteins across different group samples, and another for visualizing regulation performance (**Figure S3**). To operate this module, MS spectra and protein databases (such as those from Uniport) are uploaded to GDAS using the MSFragger-Glyco tool [1]. Biological or technical replicates are organized into different multiple input MS spectra sets, which are then used to calculate Fold-change (FC) values (**Eq. 1**).

$$FC = \frac{\sum_{i=1}^{n} x_i/n}{\sum_{i=1}^{n} x_i'/n} \qquad \text{Eq. 1}$$

The Threshold for analysis is established at the beginning of the Total Analysis Module; only FC values above this threshold are carried forward for subsequent intersection analysis. In this module, both FC values and t-test results are used to determine which proteins proceed to the visualization step. GDAS processes the uploaded large protein database to extract the significantly glycosylated proteins, creating a reduced glycoprotein database file in the required format for the next module. Crucially, the input for the next module continues to use the MS spectra raw data from diverse platforms in addition to this newly generated glycoprotein database. Finally, this process enables the estimation of volcano plots and differential expression histograms for the visualization component [2].

**The Total Analysis Module outputs data for the N/O-Glycosylation Module.**

The N/O-Glycosylation Module initiates different open-source MS data search platforms using a complete JSON (JavaScript Object Notation) file (**Figure S1** and **Figure S3**). The search utilizes a protein database containing the intersection results from the previous module. With the platform's rational settings triggered, the output is generated in various file types. GDAS uses results uploaded from platforms like GlycReSoft [3] or MetaMorpheus [4] to respectively set the disease and control groups, and a similar algorithm is employed for calculating FC values. Because glycans are central to the complexity of protein glycosylation in disease, the resulting glycosylation information takes a core position in the GDAS output, detailing the intensity of glycosylation sites and the relationship between glycan and intensity. Visualization is a critical function of GDAS, based on the glycosylation pattern extracted from the search engine output. Concurrently, glycan and glycosylation site information is exported as Excel-readable files for subsequent analysis, and a cache is established for the final analysis, containing the glycan-intensity data pairs in a special data type.

**Final Analysis Module with XGBoost and Random Forest algorithms**

The final analysis generates four key matrices: the FC/p-value expression difference matrix, the glycan intensity data pair matrix, the site-intensity data pair matrix, and the Byonic results analysis matrix (**Figure S4**). The FC/p-value matrix data originates from the total analysis, while both the glycan intensity data pair and the site intensity data pair are derived from the N/O-glycosylation module. Each of these four data categories is subjected to Bootstrap statistical methods or the Bayesian method to obtain performance difference statistics—including p-values, observations, and confidence intervals (95% CI/HDI)—which together form a statistical matrix for each glycoprotein, utilized in subsequent steps. To mitigate multicollinearity, eXtreme Gradient Boosting (XGBoost 5) [5] will be employed to build a regression model that corrects for the influence of secondary factors on the main effect. The Glycoprotein Data Analysis System (GDAS) is designed to use the fold-change matrix as the dependent variable and the other matrices as independent variables in the linear model; following model creation, a weighting matrix is used to calculate individual factor values, resulting in a score matrix. Additionally, Random Forest is used for classification, serving a function related to principal component analysis (PCA) [6] , where the glycoprotein is the dependent variable and the other matrices are the independent variables; the resulting feature correction matrix corrects the values of each statistic to form the final statistic matrix. Finally, PCA is performed using this final statistic matrix to analyze the distribution of glycoprotein taxa and visualize the output.

**Statistics methods**

**Independent two-sample t-test**

The t-test is a method of hypothesis testing whose inferences rely on several assumptions: the sample must be randomly selected, the samples must be independent of each other, the overall sample must be normally distributed, and the overall variance should be equal or approximately equal. The experimenter must often ensure the selection and randomization of the sample [7]. Furthermore, if the sample size is not sufficiently large to ensure a normal distribution, it is preferable to use Levene's chi-square test for variance [8] to determine the applicability of the t-test when dealing with smaller samples (**Eq. 2**). The entire t-test process is contained within the GDAS [9],

The Levene test for equal variances is calculated,

$$F^L = \frac{(N-k)\sum_{i=1}^{k} n_i(\bar{Z}_{i.}-\bar{Z})^2}{(k-1)\sum_{i=1}^{k}\sum_{j=1}^{n_i}(Z_{ij}-\bar{Z}_{i.})^2} \qquad \text{Eq. 2}$$

The t-test is then calculated for the means of two independent samples of scores, and the p-value is finally estimated. Based on the fold change (FC) and the p-value, the original hypothesis is either accepted or rejected [10]. The t-test is a common statistical distribution frequently used with biological samples to assess differences in the same factor between two sample groups. For expression level statistics, GDAS utilizes both the t-test and the Levene test; these are intended to discover the relationship between gene expression and disease.

**Bootstrap method for sample size over 20**

The Bootstrap method [11,12] is a statistical technique utilized in the GDAS for machine learning purposes (**Figure S5**). This method creates numerous surrogate datasets that closely resemble the original population by resampling with replacement, allowing researchers to examine the variability and uncertainty associated with their measures of interest. A key advantage of the Bootstrap method is its ability to generate a reasonably accurate sampling distribution, even with a small amount of initial data, and without relying on strict parametric assumptions about the data's distribution. The process implemented in GDAS involves: determining the number of inspection datasets, calculating group statistics, checking data validity, calculating observational effect sizes, initializing a parallel computing framework and creating parallel tasks, collecting and filtering the Bootstrap results, and finally, calculating the confidence intervals and p-values [13]. A small sample size is a limitation for the bootstrap method, especially when the count is less than 20, as statistical power may be lost. To mitigate this issue, the strategy implemented in

GDAS is to apply the bootstrap method only when the sample size is equal to or exceeds 20.

**Bayesian method for sample size under 20**

The Bayesian statistical method is another approach rooted in Bayes' theorem, which is fundamentally based on conditional probability [14]. Its core principle involves updating probability estimates of parameters or events by combining existing prior statistical models with new observations (likelihood) to produce a posterior distribution (**Figure S6**). In this design, the prior distribution is set as a Cauchy distribution. Compared to other common probability models, the Cauchy distribution has stronger tail distribution characteristics, making it much more robust. Utilizing the Markov Chain Monte Carlo (MCMC) Method, the Bayesian approach is capable of describing the statistical model distribution of glycans and their sites for each glycoprotein (**Eq. 3** and **4**).

$$P(A|B) = \frac{P(B|A)}{P(B)} \cdot P(A) \qquad \text{Eq. 3}$$

$$f(x; x_0, \gamma) = \frac{1}{\pi\gamma\left[1+\left(\frac{x-x_0}{\gamma}\right)^2\right]} = \frac{1}{\pi}\left[\frac{\gamma}{(x-x_0)^2+\gamma^2}\right] \qquad \text{Eq. 4}$$

There are two basic assumptions for these calculations: First, the distribution of glycans and sites must be consistent with the principles of Markov chains. Second, the distribution of glycans and sites must be non-periodic, meaning the probability distribution is not cyclical.

**Composition of input of overall**

The GDAS cannot directly generate final prediction results without a crucial pre-processing procedure within its architecture (**Figure S7**). The system is fundamentally divided into two loosely coupled parts—the Protein Screening and Analysis Module (comprising the Reload and Analysis Modules) and the Machine Learning Module (the Final Module)—communicating solely through data transmission managed by a Graphical User Interface (GUI). The Analysis Module first filters detected proteins using user-defined thresholds and performs initial statistical analysis (like volcano plots), while the subsequent Machine Learning Module is responsible for data reading, pre-processing, feature engineering, and integration before feeding the unified data into the XGBoost algorithm. Since statistical outputs introduce diverse data types (e.g., Excel files, lists, and tuples), and the modular design, while promoting flexibility, leads to low coupling between the modules, a necessary strategy was adopted: the data types are unified immediately after file reading to ensure consistency for all subsequent analytical steps, addressing the difficulty of entirely preventing the low-coupling issue.

**Machine Learning Algorithm**

**Linear regression and XGBoost**

While a linear regression model derived from an experiment strictly adhering to the principles of the ideal control variable method would theoretically be an exact single independent variable function with the Residual Sum of Squares (RSS) equal to zero (**Eq. 5** and **Eq. 6**), this outcome represents an idealized theoretical limit under perfect experimental control. In practical, real-world scenarios, achieving RSS = 0 is virtually impossible, as it would imply a perfect, deterministic relationship between the variables with absolutely no residual error or noise.

$$Y = kx + b \qquad \text{Eq. 5}$$

$$RSS = \sum_{i=1}^{n}(Y_i - \hat{Y}_p)^2 = 0 \qquad \text{Eq. 6}$$

In reality, the linear model used to express a system where a single variable is being controlled often involves functions with multiple independent variables (Eq. 7) [15].

$$Y = X_0 + A_1X_1 + A_2X_2 + A_3X_3 + A_4X_4 + \cdots + A_nX_n \quad \text{Eq. 7}$$

Fitting a function that closely approximates the true underlying model becomes significantly more difficult when dealing with a small number of samples (**Figure S8**). While penalty functions are often employed to adjust linear models in such cases, many experimental situations are better described by nonlinear models (like those based on differential equations) than linear ones. The decision tree is a highly popular nonlinear model known for its strong performance in weak integration learning, but its main challenge is overfitting. To overcome this challenge, the holistic approach of iteratively combining multiple weak learners to form a reliable strong learner was adopted. This concept is integrated and encapsulated by eXtreme Gradient Boosting (XGBoost), which is a highly optimized implementation of the Gradient Boosting Decision Tree (GBDT) framework (**Figure S8**). This robust algorithm will be used in the present design to fit a nonlinear model to the setup eigenvalues, and GDAS incorporates this algorithm to provide a final, robust prediction of biomarkers.

**Classification and regression using Random Forest.**

Random Forest is an ensemble machine learning algorithm developed to address the limitations of individual decision trees, which are commonly used for multi-factor analyses in biomarker discovery [16]. While a single decision tree is sensitive to default values and prone to overfitting when many factors are influential or correlated, Random Forest mitigates these drawbacks by leveraging the bagging (Bootstrap Aggregation) concept to create a collection of integrated trees, making it a stronger learner. Its random sampling methodology is consistent with the Bootstrap method, allowing the architecture of its multiple decision trees to maximize the representation of high-dimensional data's influence on the main effect. Moreover, compared to the XGBoost algorithm, the Random Forest component in GDAS is specifically designed for classification; when combined with PCA, this classification of glycoproteins can, to some extent, explain the results of KEGG pathway enrichment and protein-protein interaction analysis.

**The logic of prediction**

**Feature extraction and engineering**

The identification of a compelling biomarker hinges critically upon a robust fold-change and p-value data pair. Glycosylation plays a unique and essential role in the context of protein biomarkers, notably by influencing the cell-cell interaction. Furthermore, the inherent complexity of the glycan chain significantly dictates the overall structural complexity of glycosylation at the protein level. To precisely elucidate the structural features and abundance of protein-level glycosylation, enriched glycoproteins were analyzed using high-resolution mass spectrometry (HRMS). HRMS, when coupled with a specific library search program, is capable of yielding comprehensive data, including information on peptide modifications, protein glycosylation sites, and glycan chain composition.

Three distinct features were defined, corresponding to the three dimensions of interest: glycan structure, protein glycosylation sites, and differential expression. The initial definition of features for the glycan structure and glycosylation site dimensions was derived from the results of the commercial software such as Byonic or Byos [17]. For data analysis, Bayesian and Bootstrap methods were employed, and their resulting statistical outputs were used as eigenvalues for the XGBoost model fitting operation. In contrast to the statistically processed glycan and site data, the results for differential expression were not pre-processed but were instead directly cached in a file for subsequent program reading and processing (**Figure S3**). A key processing step involved aggregating all site statistics associated with a single glycoprotein into a single summary statistic; a similar approach was applied to the glycan chain data. By utilizing this pre-processed statistical data, the XGBoost model is expected to effectively characterize the influence of specific glycosylation features on a protein's overall differential expression.

**Statistical analysis and predictive modelling of glycosylation**

For each glycan and site associated with a glycoprotein, the Bayesian statistical analysis yields three key outputs: an observation (e.g., expression level), a Highest Density Interval (HDI), and a p-value. To characterize the composite data for a single protein, all observations pertaining to that protein are summarized by calculating a battery of descriptive statistics, including the mean, median, standard deviation (SD), range, interquartile range (IQR), coefficient of variation (CV), ratio of extreme deviation to median, and the Jensen-Shannon divergence. The individual p-values are combined using Fisher's Combined Probability Test [18]. Similarly, the 95% Confidence Interval (CI) or HDI is divided into upper and lower limits, for which the mean, SD, and range width are calculated. Initial exploration using a simple linear model, specifically XGBoost regression, demonstrated a poor fit, with the resulting function variables explaining only 0.001% of the variance. This result, along with further mathematical operations on the features, confirmed the inherently non-linear relationship among the features.

Once the model is generated, all calculated statistical values are input into the models to compute the final impact score. The regression model built using XGBoost essentially standardizes and normalizes all involved factors, ensuring that the parameters within the same system possess comparable standardized characteristics. Statistically, it is only valid to compare observations of all parameters when they reside within the same standardized evaluation system. Furthermore, in the context of multiple replicated experiments, a non-linear modelling system utilizing gradient regression substantially mitigates the issue of multicollinearity. Therefore, within this relatively rational system, it is appropriate and statistically sound to use the standardized parameters for the prediction and comparison of observations. The final impact score, calculated by the model, is used to quantitatively describe the effect of the glycoprotein on the defined features.

**Mass spectrometric analysis of intact glycopeptides**

The peptide sample (approximately 1-2 μg) was introduced onto a C18 AQ resin column and separated at a flow rate of 400 μL/min using a linear gradient (as detailed in **Table S3**). The eluting peptides were ionized by electrospray ionization (ESI) at 2.6 kV and analyzed in a data-dependent acquisition (DDA) mode with a 3 s duty cycle in positive ion mode. Full-scan mass spectra (MS1) were acquired over m/z 350 - 1800 at a resolution of 120,000, with a maximum ion accumulation time of 50 ms and AGC set to $1 \times 10^6$. Precursor ions with intensity above $2.5 \times 10^4$ were selected for MS2 acquisition, utilizing a 30 s dynamic exclusion, AGC $5 \times 10^4$, and prioritizing the highest charge state. MS/MS spectra were generated by high-energy collisional dissociation (HCD) with a 2 m/z isolation width (excluding singly-charged and unassigned ions), acquired over m/z 120 - 3000 at a resolution of 15,000 (at m/z 200), and a maximum ion accumulation time of 22 ms; for glycopeptide analysis, a stepped collision energy of 15%, 30%, and 45% was employed, and the mass list was restricted by **Table S4**, with a 1% underfill ratio.

9. Efron, B. & Hastie, T. Computer Age Statistical Inference: Algorithms, Evidence, and

Data Science. (Cambridge University Press, Cambridge; 2016).

18. Yu, X., Li, D. & Xue, L. Fisher's combined probability test for high-dimensional

covariance matrices. *J. Am. Stat. Assoc.* **119**, 511-524 (2024).

19. Martin, G.P., Riley, R.D., Collins, G.S. & Matthew, S. Developing clinical prediction models when adhering to minimum sample size recommendations: The importance of quantifying bootstrap variability in tuning parameters and predictive performance. *Stat. Methods Med. Res.* **30**, 2545-2561 (2021).

20. Zhang, Z. Univariate description and bivariate statistical inference: the first step delving into data. *Ann. Transl. Med.* **4**, 1-7 (2016).

**Supporting Tables and Figures**

**Table S1. The glycan databased used in intact glycopeptide analysis by GDAS.** (A) N-glycan database; (B) O-glycan database.

(A)

| **Glycan composition** |
|---|
| HexNAc(2)Hex(3) |
| HexNAc(2)Hex(3)Fuc(1) |
| HexNAc(2)Hex(4) |
| HexNAc(3)Hex(3) |
| HexNAc(2)Hex(3)Fuc(2) |
| HexNAc(2)Hex(4)Fuc(1) |
| HexNAc(2)Hex(5) |
| HexNAc(3)Hex(3)Fuc(1) |
| HexNAc(3)Hex(4) |
| HexNAc(4)Hex(3) |
| HexNAc(2)Hex(5)Fuc(1) |
| HexNAc(2)Hex(6) |
| HexNAc(3)Hex(3)Fuc(2) |
| HexNAc(3)Hex(4)Fuc(1) |
| HexNAc(3)Hex(5) |
| HexNAc(4)Hex(3)Fuc(1) |
| HexNAc(4)Hex(4) |
| HexNAc(5)Hex(3) |
| HexNAc(2)Hex(7) |

HexNAc(3)Hex(4)NeuAc(1)

HexNAc(3)Hex(4)Fuc(2)

HexNAc(3)Hex(4)NeuGc(1)

HexNAc(3)Hex(5)Fuc(1)

HexNAc(3)Hex(6)

HexNAc(4)Hex(3)NeuAc(1)

HexNAc(4)Hex(3)Fuc(2)

HexNAc(4)Hex(3)NeuGc(1)

HexNAc(4)Hex(4)Fuc(1)

HexNAc(4)Hex(5)

HexNAc(5)Hex(3)Fuc(1)

HexNAc(5)Hex(4)

HexNAc(3)Hex(4)Fuc(1)NeuAc(1)

HexNAc(2)Hex(8)

HexNAc(6)Hex(3)

HexNAc(3)Hex(4)Fuc(1)NeuGc(1)

HexNAc(3)Hex(5)NeuAc(1)

HexNAc(3)Hex(5)NeuGc(1)

HexNAc(3)Hex(6)Fuc(1)

HexNAc(4)Hex(3)Fuc(3)

HexNAc(4)Hex(4)NeuAc(1)

HexNAc(4)Hex(4)Fuc(2)

HexNAc(4)Hex(4)NeuGc(1)

HexNAc(4)Hex(5)Fuc(1)

HexNAc(4)Hex(6)

HexNAc(5)Hex(4)Fuc(1)

HexNAc(5)Hex(5)

HexNAc(6)Hex(3)Fuc(1)

HexNAc(2)Hex(9)

HexNAc(6)Hex(4)

HexNAc(3)Hex(6)NeuAc(1)

HexNAc(3)Hex(6)NeuGc(1)

HexNAc(4)Hex(4)Fuc(1)NeuAc(1)

HexNAc(7)Hex(3)

HexNAc(4)Hex(4)Fuc(1)NeuGc(1)

HexNAc(4)Hex(5)NeuAc(1)

HexNAc(4)Hex(5)Fuc(2)

HexNAc(4)Hex(5)NeuGc(1)

HexNAc(4)Hex(6)Fuc(1)

HexNAc(5)Hex(3)Fuc(1)NeuAc(1)

HexNAc(4)Hex(7)

HexNAc(5)Hex(4)NeuAc(1)

HexNAc(5)Hex(3)Fuc(1)NeuGc(1)

HexNAc(5)Hex(4)Fuc(2)

HexNAc(5)Hex(4)NeuGc(1)

HexNAc(5)Hex(5)Fuc(1)

HexNAc(5)Hex(6)

HexNAc(6)Hex(3)Fuc(2)

HexNAc(6)Hex(4)Fuc(1)

HexNAc(3)Hex(6)Fuc(1)NeuAc(1)

HexNAc(2)Hex(10)

HexNAc(6)Hex(5)

HexNAc(3)Hex(6)Fuc(1)NeuGc(1)

HexNAc(4)Hex(4)Fuc(2)NeuAc(1)

HexNAc(7)Hex(3)Fuc(1)

HexNAc(4)Hex(5)Fuc(1)NeuAc(1)

HexNAc(4)Hex(4)Fuc(2)NeuGc(1)

HexNAc(4)Hex(5)Fuc(3)

HexNAc(7)Hex(4)

HexNAc(4)Hex(6)NeuAc(1)

HexNAc(4)Hex(5)Fuc(1)NeuGc(1)

HexNAc(4)Hex(6)Fuc(2)

HexNAc(4)Hex(6)NeuGc(1)

HexNAc(4)Hex(7)Fuc(1)

HexNAc(5)Hex(4)Fuc(1)NeuAc(1)

HexNAc(8)Hex(3)

HexNAc(5)Hex(4)Fuc(1)NeuGc(1)

HexNAc(5)Hex(5)NeuAc(1)

HexNAc(5)Hex(5)Fuc(2)

HexNAc(5)Hex(5)NeuGc(1)

HexNAc(5)Hex(6)Fuc(1)

HexNAc(6)Hex(3)Fuc(1)NeuAc(1)

HexNAc(6)Hex(3)Fuc(3)

HexNAc(5)Hex(7)

HexNAc(6)Hex(4)NeuAc(1)

HexNAc(6)Hex(3)Fuc(1)NeuGc(1)

HexNAc(6)Hex(4)Fuc(2)

HexNAc(6)Hex(5)Fuc(1)

HexNAc(2)Hex(11)

HexNAc(6)Hex(6)

HexNAc(4)Hex(5)NeuAc(1)

HexNAc(4)Hex(5)NeuAc(2)

HexNAc(4)Hex(5)Fuc(2)NeuAc(1)

HexNAc(4)Hex(5)Fuc(4)

HexNAc(7)Hex(4)Fuc(1)

HexNAc(4)Hex(5)NeuAc(1)NeuGc(1)

HexNAc(4)Hex(5)Fuc(2)NeuGc(1)

HexNAc(4)Hex(6)Fuc(1)NeuAc(1)

HexNAc(4)Hex(6)Fuc(3)

HexNAc(4)Hex(5)NeuGc(2)

HexNAc(4)Hex(6)Fuc(1)NeuGc(1)

HexNAc(4)Hex(7)NeuAc(1)

HexNAc(4)Hex(7)Fuc(2)

HexNAc(5)Hex(4)NeuAc(1)

HexNAc(5)Hex(4)NeuAc(2)

HexNAc(5)Hex(4)Fuc(2)NeuAc(1)

HexNAc(8)Hex(3)Fuc(1)

HexNAc(5)Hex(4)NeuAc(1)NeuGc(1)

HexNAc(5)Hex(5)Fuc(1)NeuAc(1)

HexNAc(5)Hex(4)Fuc(2)NeuGc(1)

HexNAc(5)Hex(5)Fuc(3)

HexNAc(8)Hex(4)

HexNAc(5)Hex(4)NeuGc(2)

HexNAc(5)Hex(5)Fuc(1)NeuGc(1)

HexNAc(5)Hex(6)NeuAc(1)

HexNAc(5)Hex(6)Fuc(2)

HexNAc(6)Hex(3)Fuc(2)NeuAc(1)

HexNAc(5)Hex(6)NeuGc(1)

HexNAc(5)Hex(7)Fuc(1)

HexNAc(6)Hex(3)Fuc(2)NeuGc(1)

HexNAc(5)Hex(8)

HexNAc(9)Hex(3)

HexNAc(6)Hex(5)Fuc(2)

HexNAc(6)Hex(6)Fuc(1)

HexNAc(2)Hex(12)

HexNAc(4)Hex(5)Fuc(1)NeuAc(2)

HexNAc(4)Hex(5)Fuc(1)NeuAc(1)

HexNAc(6)Hex(7)

HexNAc(7)Hex(4)Fuc(2)

HexNAc(4)Hex(5)Fuc(1)NeuAc(1)NeuGc(1)

HexNAc(4)Hex(5)Fuc(1)NeuGc(2)

HexNAc(5)Hex(4)Fuc(1)NeuAc(2)

HexNAc(5)Hex(4)Fuc(1)NeuAc(1)

HexNAc(7)Hex(6)

HexNAc(5)Hex(5)NeuAc(2)

HexNAc(5)Hex(5)NeuAc(1)

HexNAc(5)Hex(4)Fuc(1)NeuAc(1)NeuGc(1)

HexNAc(5)Hex(4)Fuc(1)NeuGc(2)

HexNAc(5)Hex(5)NeuAc(1)NeuGc(1)

HexNAc(5)Hex(6)Fuc(1)NeuAc(1)

HexNAc(5)Hex(6)Fuc(3)

HexNAc(6)Hex(3)Fuc(1)NeuAc(1)

HexNAc(6)Hex(3)Fuc(1)NeuAc(2)

HexNAc(8)Hex(5)

HexNAc(5)Hex(5)NeuGc(2)

HexNAc(5)Hex(6)Fuc(1)NeuGc(1)

HexNAc(6)Hex(3)Fuc(1)NeuAc(1)NeuGc(1)

HexNAc(5)Hex(8)Fuc(1)

HexNAc(9)Hex(3)Fuc(1)

HexNAc(6)Hex(3)Fuc(1)NeuGc(2)

HexNAc(9)Hex(4)

HexNAc(6)Hex(6)Fuc(2)

HexNAc(6)Hex(7)Fuc(1)

HexNAc(7)Hex(6)Fuc(1)

HexNAc(5)Hex(5)Fuc(1)NeuAc(2)

HexNAc(5)Hex(5)Fuc(1)NeuAc(1)

HexNAc(7)Hex(7)

HexNAc(5)Hex(5)Fuc(1)NeuAc(1)NeuGc(1)

HexNAc(5)Hex(6)NeuAc(1)

HexNAc(5)Hex(6)NeuAc(2)

HexNAc(5)Hex(6)Fuc(2)NeuAc(1)

HexNAc(8)Hex(5)Fuc(1)

HexNAc(5)Hex(5)Fuc(1)NeuGc(2)

HexNAc(5)Hex(6)NeuAc(1)NeuGc(1)

HexNAc(5)Hex(7)Fuc(1)NeuAc(1)

HexNAc(5)Hex(6)Fuc(2)NeuGc(1)

HexNAc(8)Hex(6)

HexNAc(5)Hex(6)NeuGc(2)

HexNAc(5)Hex(9)Fuc(1)

HexNAc(9)Hex(4)Fuc(1)

HexNAc(6)Hex(6)Fuc(3)

HexNAc(4)Hex(5)Fuc(3)NeuAc(2)

HexNAc(4)Hex(5)Fuc(3)NeuAc(1)

HexNAc(6)Hex(7)NeuAc(1)

HexNAc(4)Hex(5)Fuc(3)NeuAc(1)NeuGc(1)

HexNAc(6)Hex(7)NeuGc(1)

HexNAc(4)Hex(5)Fuc(3)NeuGc(2)

HexNAc(6)Hex(9)

HexNAc(7)Hex(7)Fuc(1)

HexNAc(5)Hex(6)Fuc(1)NeuAc(1)

HexNAc(5)Hex(6)Fuc(1)NeuAc(2)

HexNAc(5)Hex(6)Fuc(3)NeuAc(1)

HexNAc(7)Hex(8)

HexNAc(5)Hex(6)Fuc(1)NeuAc(1)NeuGc(1)

HexNAc(5)Hex(6)Fuc(3)NeuGc(1)

HexNAc(5)Hex(6)Fuc(1)NeuGc(2)

HexNAc(8)Hex(7)

HexNAc(6)Hex(6)NeuAc(1)

HexNAc(6)Hex(6)NeuAc(2)

HexNAc(6)Hex(6)NeuAc(1)NeuGc(1)

HexNAc(6)Hex(7)Fuc(1)NeuAc(1)

HexNAc(9)Hex(6)

HexNAc(6)Hex(6)NeuGc(2)

HexNAc(6)Hex(7)Fuc(1)NeuGc(1)

HexNAc(6)Hex(8)NeuAc(1)

HexNAc(5)Hex(6)NeuAc(3)

HexNAc(5)Hex(6)NeuAc(1)

HexNAc(5)Hex(6)NeuAc(2)

HexNAc(7)Hex(8)Fuc(1)

HexNAc(5)Hex(6)NeuAc(1)NeuGc(1)

HexNAc(5)Hex(6)NeuAc(2)NeuGc(1)

HexNAc(5)Hex(7)Fuc(1)NeuAc(1)

HexNAc(5)Hex(7)Fuc(1)NeuAc(2)

HexNAc(5)Hex(6)NeuAc(1)NeuGc(2)

HexNAc(5)Hex(7)Fuc(1)NeuAc(1)NeuGc(1)

HexNAc(5)Hex(8)Fuc(4)

HexNAc(5)Hex(6)NeuGc(3)

HexNAc(5)Hex(7)Fuc(1)NeuGc(2)

HexNAc(6)Hex(6)Fuc(1)NeuAc(2)

HexNAc(6)Hex(6)Fuc(1)NeuAc(1)

HexNAc(8)Hex(8)

HexNAc(6)Hex(6)Fuc(1)NeuAc(1)NeuGc(1)

HexNAc(6)Hex(7)NeuAc(1)

HexNAc(6)Hex(7)NeuAc(2)

HexNAc(6)Hex(7)Fuc(4)

HexNAc(9)Hex(6)Fuc(1)

HexNAc(6)Hex(7)NeuAc(1)NeuGc(1)

HexNAc(6)Hex(6)Fuc(1)NeuGc(2)

HexNAc(6)Hex(8)Fuc(1)NeuAc(1)

HexNAc(6)Hex(7)NeuGc(2)

HexNAc(5)Hex(6)Fuc(1)NeuAc(3)

HexNAc(5)Hex(6)Fuc(1)NeuAc(1)

HexNAc(5)Hex(6)Fuc(1)NeuAc(2)

HexNAc(7)Hex(8)NeuAc(1)

HexNAc(5)Hex(6)Fuc(1)NeuAc(2)NeuGc(1)

HexNAc(5)Hex(6)Fuc(1)NeuAc(1)NeuGc(1)

HexNAc(7)Hex(8)NeuGc(1)

HexNAc(5)Hex(6)Fuc(1)NeuAc(1)NeuGc(2)

HexNAc(6)Hex(5)Fuc(1)NeuAc(2)

HexNAc(6)Hex(5)Fuc(1)NeuAc(1)

HexNAc(6)Hex(5)Fuc(1)NeuAc(3)

HexNAc(5)Hex(6)Fuc(1)NeuGc(3)

HexNAc(6)Hex(5)Fuc(1)NeuAc(1)NeuGc(1)

HexNAc(6)Hex(5)Fuc(1)NeuAc(2)NeuGc(1)

HexNAc(8)Hex(8)Fuc(1)

HexNAc(6)Hex(5)Fuc(1)NeuAc(1)NeuGc(2)

HexNAc(6)Hex(7)Fuc(1)NeuAc(2)

HexNAc(8)Hex(9)

HexNAc(6)Hex(7)Fuc(5)

HexNAc(6)Hex(5)Fuc(1)NeuGc(3)

HexNAc(6)Hex(11)Fuc(1)

HexNAc(10)Hex(7)

HexNAc(6)Hex(6)Fuc(1)NeuAc(2)

HexNAc(6)Hex(6)Fuc(1)NeuAc(3)

HexNAc(6)Hex(6)Fuc(1)NeuAc(2)NeuGc(1)

HexNAc(6)Hex(7)NeuAc(3)

HexNAc(6)Hex(7)Fuc(4)NeuAc(1)

HexNAc(8)Hex(9)Fuc(1)

HexNAc(6)Hex(7)NeuAc(2)NeuGc(1)

HexNAc(6)Hex(6)Fuc(1)NeuAc(1)NeuGc(2)

HexNAc(6)Hex(7)Fuc(4)NeuGc(1)

HexNAc(6)Hex(6)Fuc(1)NeuGc(3)

HexNAc(6)Hex(10)Fuc(1)NeuAc(1)

HexNAc(7)Hex(7)Fuc(1)NeuAc(2)

HexNAc(6)Hex(10)Fuc(1)NeuGc(1)

HexNAc(6)Hex(7)Fuc(1)NeuAc(3)

HexNAc(6)Hex(9)Fuc(1)NeuAc(2)

HexNAc(6)Hex(9)Fuc(1)NeuAc(1)NeuGc(1)

HexNAc(9)Hex(9)Fuc(1)

HexNAc(7)Hex(8)Fuc(1)NeuAc(1)

HexNAc(7)Hex(8)Fuc(1)NeuAc(2)

HexNAc(9)Hex(10)

HexNAc(7)Hex(8)Fuc(1)NeuAc(1)NeuGc(1)

HexNAc(7)Hex(8)Fuc(1)NeuGc(2)

HexNAc(6)Hex(7)NeuAc(4)

HexNAc(6)Hex(7)NeuAc(3)NeuGc(1)

HexNAc(6)Hex(7)NeuAc(2)NeuGc(2)

HexNAc(7)Hex(7)Fuc(1)NeuAc(3)

HexNAc(7)Hex(7)Fuc(1)NeuAc(2)NeuGc(1)

HexNAc(9)Hex(10)Fuc(1)

HexNAc(7)Hex(7)Fuc(1)NeuAc(1)NeuGc(2)

HexNAc(6)Hex(7)Fuc(1)NeuAc(4)

HexNAc(6)Hex(7)Fuc(1)NeuAc(3)NeuGc(1)

HexNAc(7)Hex(6)Fuc(1)NeuAc(4)

HexNAc(7)Hex(6)Fuc(1)NeuAc(3)NeuGc(1)

HexNAc(7)Hex(8)Fuc(1)NeuAc(1)

HexNAc(7)Hex(8)Fuc(1)NeuAc(3)

HexNAc(7)Hex(8)Fuc(1)NeuAc(2)

HexNAc(7)Hex(8)Fuc(1)NeuAc(2)NeuGc(1)

HexNAc(7)Hex(8)Fuc(1)NeuAc(1)NeuGc(1)

HexNAc(7)Hex(8)Fuc(1)NeuAc(1)NeuGc(2)

HexNAc(7)Hex(8)Fuc(1)NeuGc(3)

HexNAc(10)Hex(10)Fuc(1)

HexNAc(7)Hex(7)Fuc(1)NeuAc(4)

HexNAc(7)Hex(8)Fuc(1)NeuAc(4)

| |
|---|
| HexNAc(8)Hex(9)Fuc(1)NeuAc(3) |
| HexNAc(8)Hex(9)Fuc(1)NeuAc(2)NeuGc(1) |
| HexNAc(8)Hex(9)Fuc(1)NeuAc(1)NeuGc(2) |
| HexNAc(11)Hex(11)NeuAc(1) |
| HexNAc(11)Hex(11)NeuGc(1) |
| HexNAc(8)Hex(9)Fuc(1)NeuAc(4) |
| HexNAc(9)Hex(10)Fuc(1)NeuAc(4) |

(B)

| **O-glycan composition** |
|---|
| HexNAc(1) |
| HexNAc(1)Hex(1) |
| HexNAc(2) |
| HexNAc(1)Hex(1)Fuc(1) |
| HexNAc(2)Hex(1) |
| HexNAc(3) |
| HexNAc(1)Hex(1)NeuAc(1) |
| HexNAc(1)Hex(1)NeuGc(1) |
| HexNAc(1)Hex(2)Fuc(1) |
| HexNAc(2)Hex(1)Fuc(1) |
| HexNAc(2)Hex(2) |
| HexNAc(3)Fuc(1) |
| HexNAc(3)Hex(1) |
| HexNAc(1)Hex(1)Fuc(1)NeuAc(1) |
| HexNAc(1)Hex(1)Fuc(1)NeuGc(1) |

HexNAc(2)Hex(1)NeuAc(1)

HexNAc(2)Hex(1)Fuc(2)

HexNAc(2)Hex(1)NeuGc(1)

HexNAc(2)Hex(2)Fuc(1)

HexNAc(2)Hex(3)

HexNAc(3)Hex(1)Fuc(1)

HexNAc(3)Hex(2)

HexNAc(1)Hex(1)NeuAc(2)

HexNAc(1)Hex(1)NeuAc(1)NeuGc(1)

HexNAc(4)Hex(1)

HexNAc(1)Hex(1)NeuGc(2)

HexNAc(2)Hex(1)Fuc(1)NeuAc(1)

HexNAc(2)Hex(1)Fuc(1)NeuGc(1)

HexNAc(2)Hex(2)NeuAc(1)

HexNAc(2)Hex(2)Fuc(2)

HexNAc(2)Hex(3)Fuc(1)

HexNAc(3)Hex(1)NeuAc(1)

HexNAc(3)Hex(1)Fuc(2)

HexNAc(3)Hex(2)Fuc(1)

HexNAc(3)Hex(3)

HexNAc(4)Hex(1)Fuc(1)

HexNAc(4)Hex(2)

HexNAc(2)Hex(1)NeuAc(2)

HexNAc(2)Hex(2)Fuc(1)NeuAc(1)

HexNAc(2)Hex(1)Fuc(2)NeuGc(1)

HexNAc(2)Hex(2)Fuc(3)

HexNAc(2)Hex(3)NeuAc(1)

HexNAc(3)Hex(1)Fuc(1)NeuAc(1)

HexNAc(3)Hex(1)Fuc(1)NeuGc(1)

HexNAc(3)Hex(2)NeuAc(1)

HexNAc(3)Hex(2)Fuc(2)

HexNAc(1)Hex(1)NeuAc(3)

HexNAc(3)Hex(3)Fuc(1)

HexNAc(2)Hex(2)NeuAc(2)

HexNAc(2)Hex(2)Fuc(2)NeuAc(1)

HexNAc(3)Hex(2)Fuc(1)NeuAc(1)

HexNAc(3)Hex(2)Fuc(3)

HexNAc(3)Hex(3)NeuAc(1)

HexNAc(3)Hex(3)Fuc(2)

HexNAc(4)Hex(2)Fuc(2)

HexNAc(4)Hex(3)Fuc(1)

HexNAc(2)Hex(2)Fuc(1)NeuAc(2)

HexNAc(4)Hex(4)

HexNAc(5)Hex(3)

HexNAc(3)Hex(2)NeuAc(2)

HexNAc(3)Hex(2)Fuc(2)NeuAc(1)

HexNAc(3)Hex(2)Fuc(4)

HexNAc(3)Hex(3)Fuc(1)NeuAc(1)

HexNAc(3)Hex(3)Fuc(3)

HexNAc(4)Hex(3)Fuc(2)

| |
|---|
| HexNAc(3)Hex(3)NeuAc(2) |
| HexNAc(3)Hex(3)Fuc(1)NeuAc(2) |
| HexNAc(5)Hex(5) |
| HexNAc(6)Hex(3)Fuc(1) |
| HexNAc(6)Hex(4) |
| HexNAc(4)Hex(4)Fuc(3) |
| HexNAc(5)Hex(3)Fuc(1)NeuAc(1) |
| HexNAc(3)Hex(3)Fuc(2)NeuAc(2) |
| HexNAc(4)Hex(4)Fuc(2)NeuAc(1) |
| HexNAc(6)Hex(4)Fuc(2) |
| HexNAc(4)Hex(4)Fuc(3)NeuAc(1) |
| HexNAc(5)Hex(5)Fuc(3) |
| HexNAc(6)Hex(5)Fuc(3) |

**Table S2. Typical fragmentation ions produced by different MS/MS techniques.** This table summarizes the characteristic fragmentation patterns of various MS/MS methods: HCD (higher-energy collisional dissociation) uses higher collision energy to generate primarily b and y ions for peptide sequencing, often resulting in neutral loss of glycans. ETD (electron-transfer dissociation) favors peptide backbone cleavage, yielding c and z ions while leaving the glycan largely intact, though it provides little information about the glycan structure itself. EThcD (electron-transfer/higher-energy collision dissociation) is a hybrid method combining the features of HCD and ETD. EAD (electron-activated dissociation) generates a, b, x, and y ions, alongside characteristic cross-ring fragments useful for detailed glycan structure analysis.

| Ion Type | Description | HCD | ETD | EThcD | EAD |
|---|---|---|---|---|---|
| a, b, c | N-terminal peptide backbone fragments | Primarily **b** ions (at higher energy) | Primarily **c** ions | Abundant **b** and **c** ion s | Primarily **a** and **b** ions (with EAD only) |
| x, y, z | C-terminal peptide backbone fragments | Primarily **y** ions (at higher energy) | Primarily **z** ions | Abundant **y** and **z** ion s | Primarily **y** ions |
| B-ions | Glycan-only fragments from the non-reducing end | Abundant | Few/None | Abundant (especially with higher HCD energy) | Abundant |
| Y-ions | Peptide with a truncated/intact glycan attached | Abundant (especially at lower energy, for site localization) | Abundant (intact glycan on peptide) | Abundant | Abundant (intact or partial glycan on peptide) |
| Oxonium ions | Small, diagnostic glycan fragments (e.g., m/z 204.09 for HexNAc) | Abundant, used as a trigger for other methods | Low/None | Abundant | Abundant |

| | | | | | |
|---|---|---|---|---|---|
| Cross-ring ions | Ions from cleavages within the glycan ring | Few | Very few | Very few | Abundant, useful for detailed linkage analysis |

**Table S3. Nanoflow liquid chromatography gradient.** ACN = acetonitrile.

| Time (min) | Flow (μl/min) | HPLC Water (v/v, %) | ACN (v/v, %) |
|---|---|---|---|
| | 0.400 | 99.0 | 1.0 |
| 1.00 | 0.400 | 95.0 | 5.0 |
| 75.00 | 0.400 | 78.0 | 22.0 |
| 95.00 | 0.400 | 64.0 | 36.0 |
| 100.00 | 0.400 | 50.0 | 50.0 |
| 105.00 | 0.400 | 10.0 | 90.0 |
| 115.00 | 0.400 | 10.0 | 90.0 |
| 115.10 | 0.400 | 99.0 | 1.0 |
| 120.00 | 0.400 | 99.0 | 1.0 |

**Table S4. The oxonium ions used in tandem mass spectrometry analysis.**

| Type | Oxonium ion | Mass (m/z) |
|---|---|---|
| N-/O-glycopeptide | HexNAc Fragment | 138.0545 |
| | HexNAc | 204.0867 |
| N-glycopeptide | Hex | 163.0601 |
| | HexHexNAc | 366.1395 |

**Figure S1. The interface and workflow of GDAS for high-throughput identification of disease-specific glycosylation.** The workflow begins with (**A**) the interface for uploading MS raw data and the protein database, followed by specialized analyses for N-glycosylation using (**B**) the N-glycosylation analysis module, integrated with (**C**) MSFragger and GlycReSoft interfaces, and O-glycosylation using (**D**) the O-glycosylation analysis module, integrated with (**E**) MSFragger and O-Pair interfaces. The final results are processed and presented via (**F**) Byonic and the final output interface.

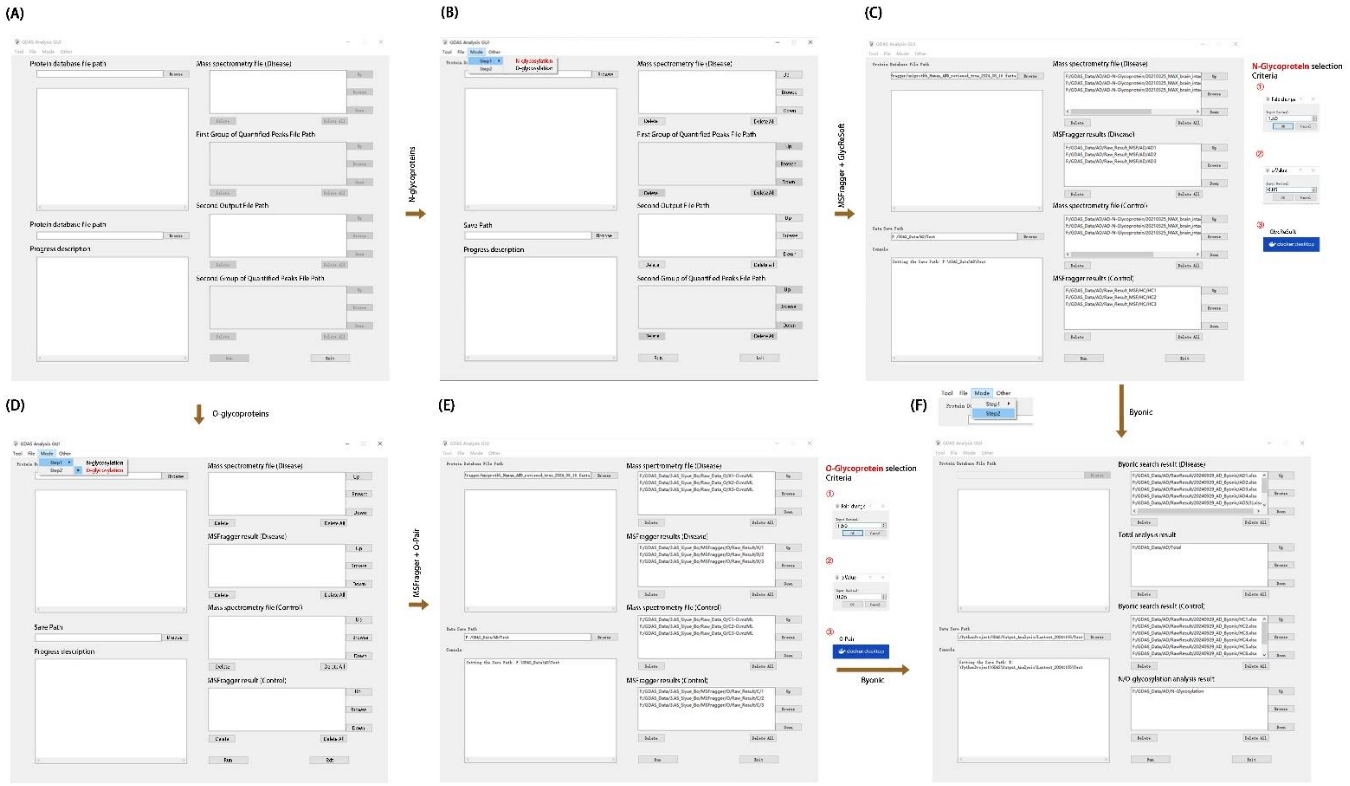

**Figure S2. The streamlined algorithm of GDAS analysis of protein glycosylation.** The large number of proteins as search database is efficiently reduced by combination of MSFragger, GlycReSoft and O-Pair. The reduced protein database allows for quick search of targeted proteins without loss of detailed site-specific glycoforms.

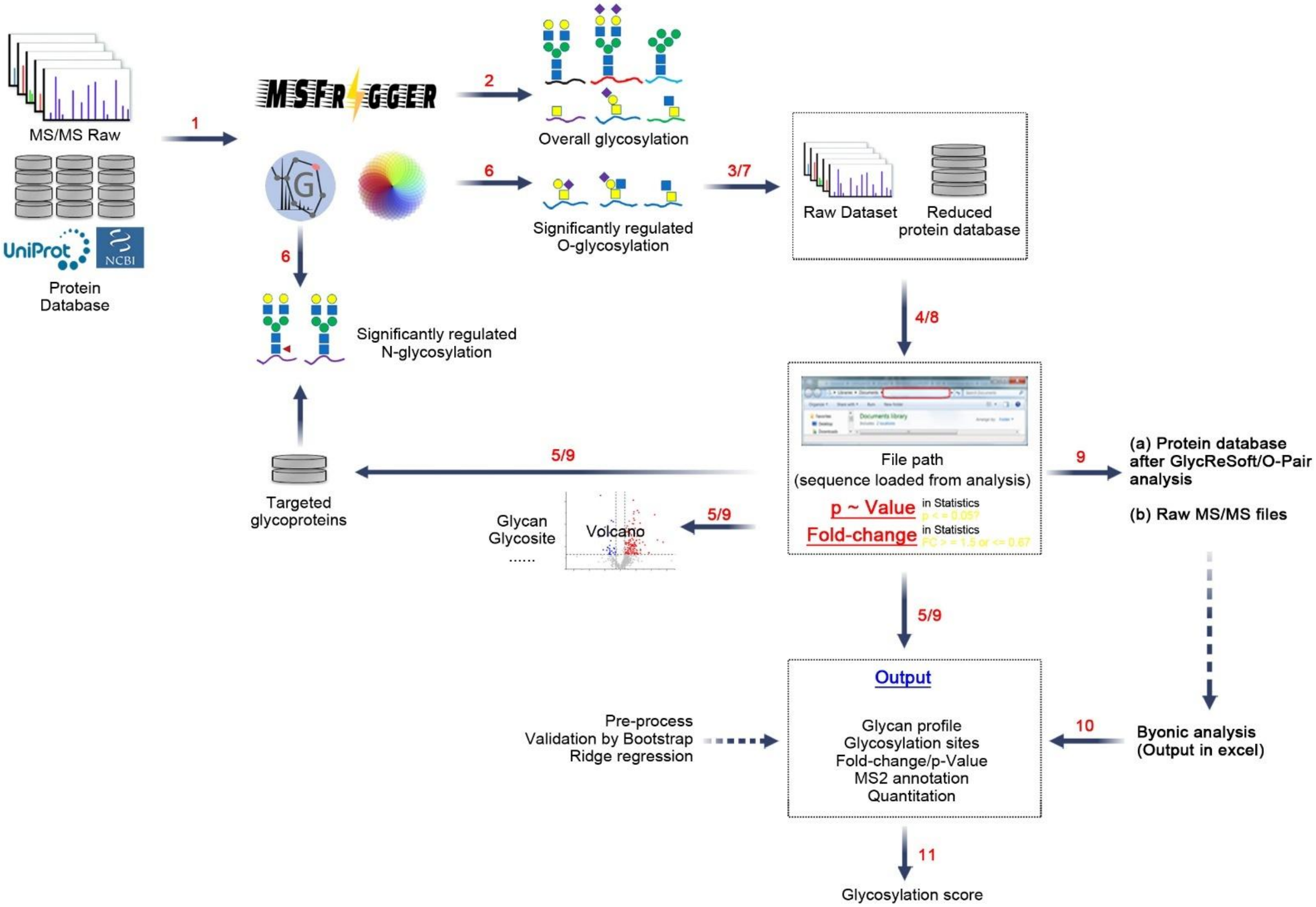

**Figure S3. GDAS algorithm procedure.** The workflow begins with (**A**) activation of the main modules, the Protein Database Reloading Module and the MSFragger/GlycReSoft/O-Pair Result Reading Module. The Result Reading Module reads the input search files, determines the set of detected proteins common to all results, and sends this intersection to the Protein Database Reloading Module. This module reads the full protein database (typically from UniProt), filters it based on the common detected proteins and applied thresholds, and outputs a new, filtered protein database. Other relevant search data are passed to the Analysis Module. (**B**) The Analysis Module uses distinct workflows based on the search result type. The reading module's output is processed to form three pairs based on the data's tree structure: Glycan-Intensity, Sites-Intensity, and Peptide-Intensity pairs. This is followed by data preprocessing, including a secondary screening of the protein library and the calculation of Fold-Change and $p$-values. For MSFragger results, a homogeneity of variance test is performed prior to Fold-Change and p-value calculation, and a volcano plot is generated. For GlycReSoft/O-Pair results, Fold-Change is calculated directly, and a Glycan Profile is summarized for each file, including GlycanMass-Intensity data and plot, as well as Site-Intensity data and plot for each glycoprotein.

(A)

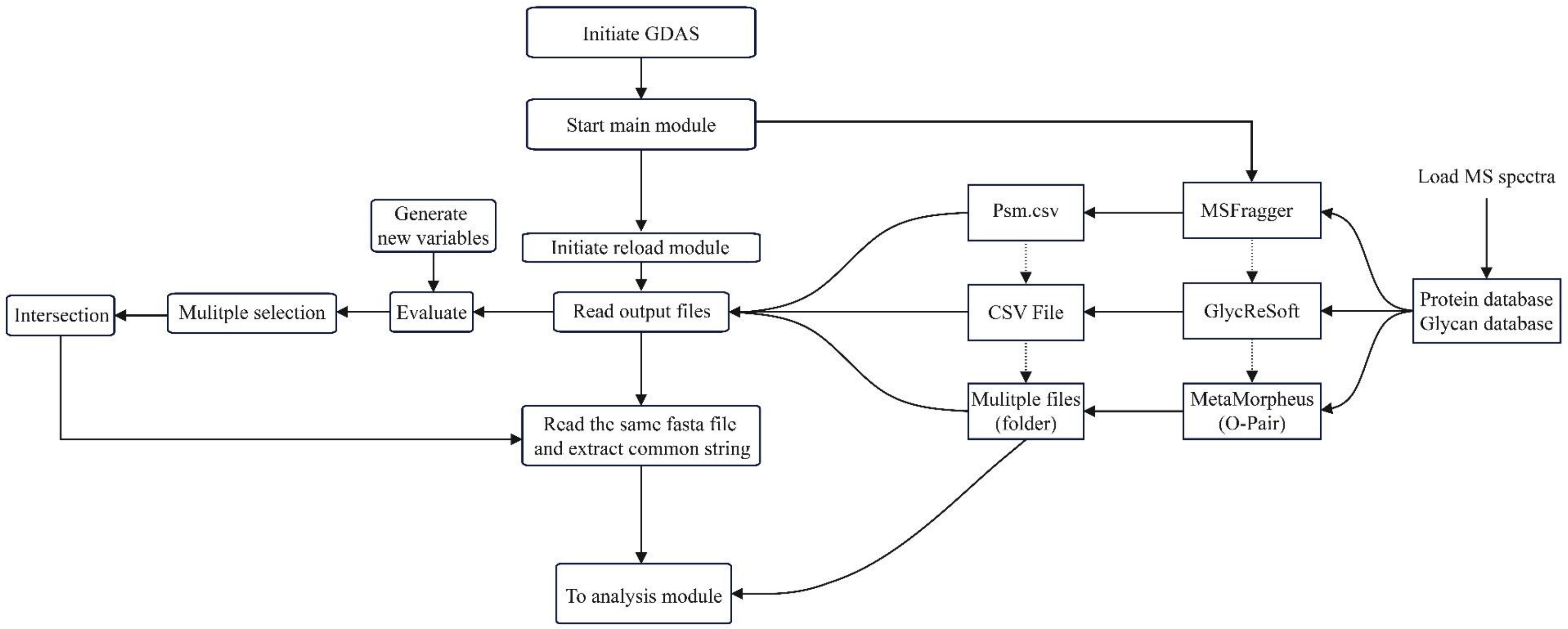

Initiate GDAS
Start main module
Initiate reload module
Generate
new variables
Read output files
Evaluate
Mulitple selection
Intersection
Read the same fasta file
and extract common string
To analysis module
Psm.csv
CSV File
Mulitple files
(folder)
MSFragger
GlycReSoft
MetaMorpheus
(O-Pair)
Load MS spectra
Protein database
Glycan database

(B)

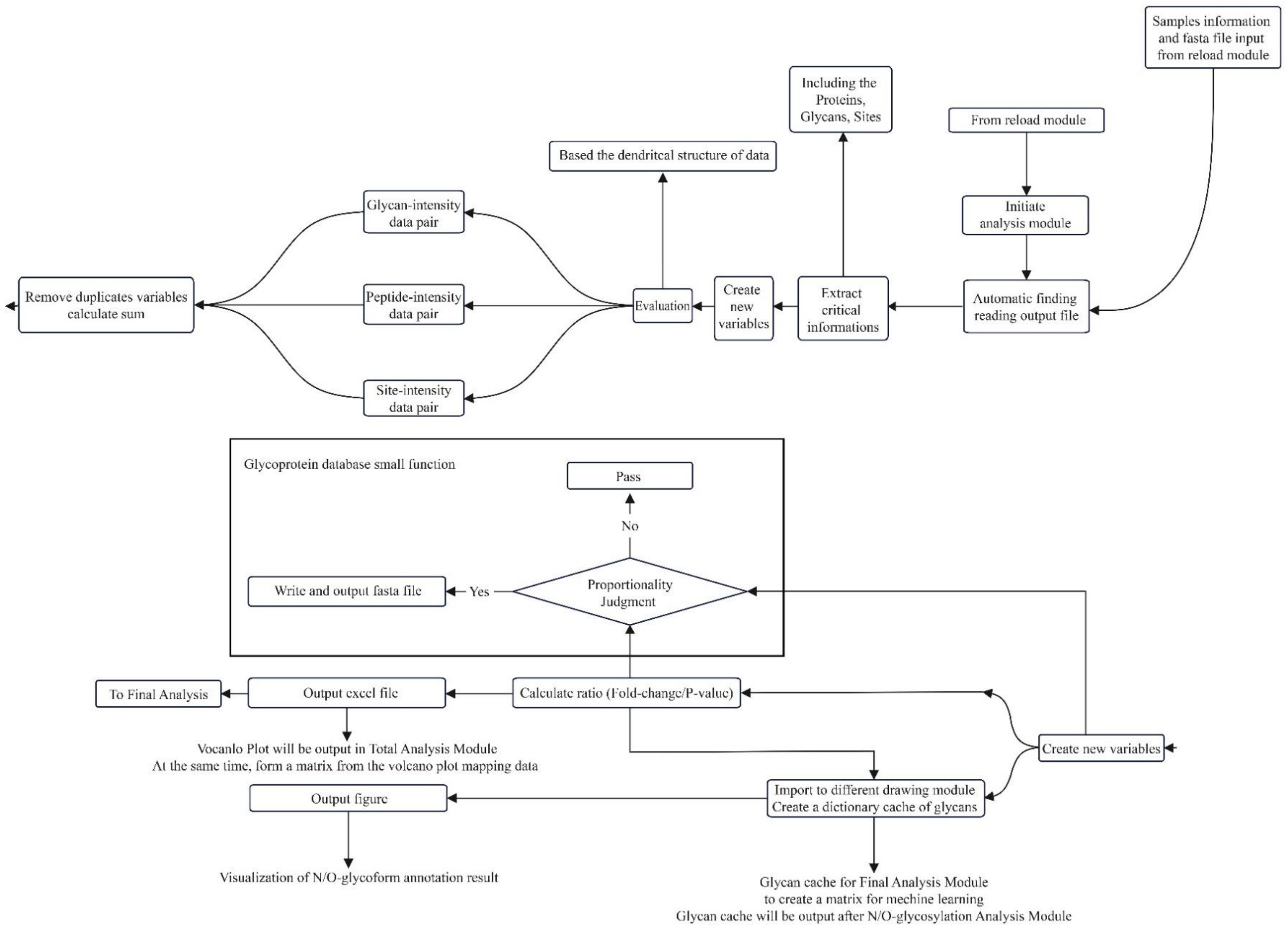

Samples information and fasta file input from reload module
Including the Proteins, Glycans, Sites
From reload module
Based the dendritcal structure of data
Initiate analysis module
Glycan-intensity data pair
Remove duplicates variables calculate sum
Peptide-intensity data pair
Evaluation
Create new variables
Extract critical informations
Automatic finding reading output file
Site-intensity data pair
Glycoprotein database small function
Pass
No
Write and output fasta file
Yes
Proportionality Judgment
To Final Analysis
Output excel file
Calculate ratio (Fold-change/P-value)
Vocanlo Plot will be output in Total Analysis Module
At the same time, form a matrix from the volcano plot mapping data
Create new variables
Output figure
Import to different drawing module
Create a dictionary cache of glycans
Visualization of N/O-glycoform annotation result
Glycan cache for Final Analysis Module
to create a matrix for mechine learning
Glycan cache will be output after N/O-glycosylation Analysis Module

**Figure S4. GDAS machine learning procedure.** The machine learning module is structured into three consecutive parts: data reading and preprocessing, feature engineering, and the machine learning process itself. Data reading involves gathering the four key input datasets required by the XGBoost algorithm: the Fold-Change/$p$-Value dataset, the Glycans-Intensity dataset, the Sites-Intensity dataset, and the Byonic results. After module initialization, these datasets are read and categorized by their number of data points for feature engineering. If a dataset contains 20 or more data points, the Bootstrap method is applied for feature engineering; otherwise, Bayesian statistical analysis is used. The Glycans-Intensity, Sites-Intensity, and Byonic results undergo this feature engineering process, while the Fold-Change/$p$-Value dataset bypasses feature engineering and is fed directly into XGBoost as independent variables. Following the fitting of the XGBoost model, the independent variables are substituted into the fitted model, and the predicted value is determined by taking the average of all predicted values obtained through grid search.

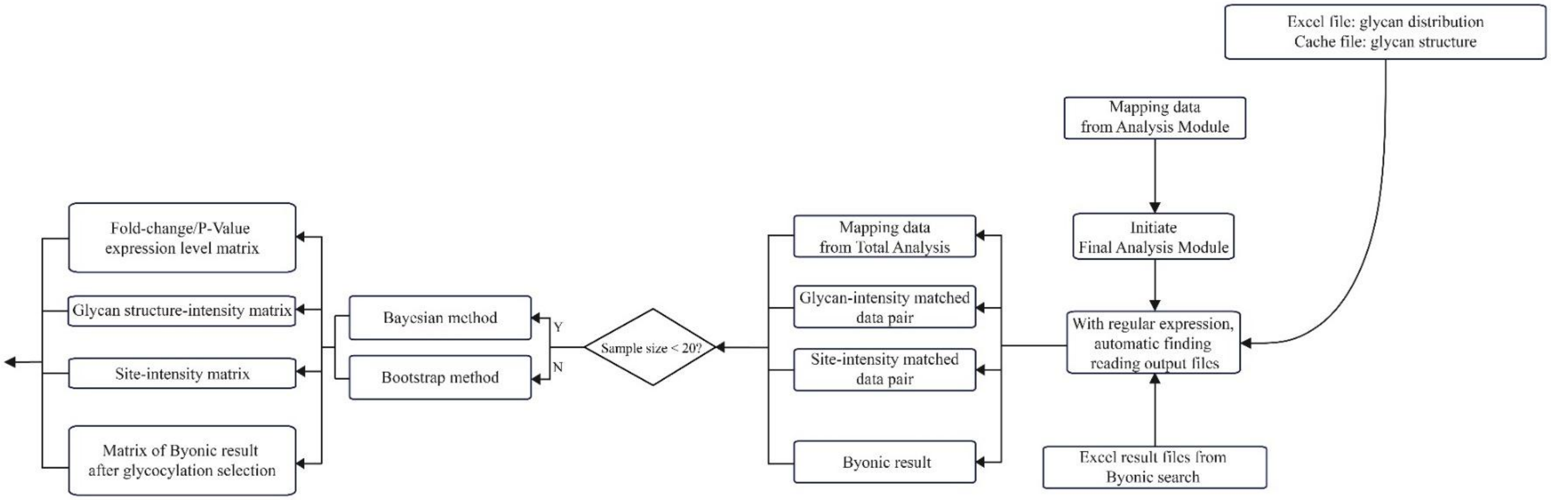
Excel file: glycan distribution
Cache file: glycan structure
Mapping data from Analysis Module
Initiate Final Analysis Module
With regular expression, automatic finding reading output files
Excel result files from Byonic search
Mapping data from Total Analysis
Glycan-intensity matched data pair
Site-intensity matched data pair
Byonic result
Sample size < 20?
Y
N
Bayesian method
Bootstrap method
Fold-change/P-Value expression level matrix
Glycan structure-intensity matrix
Site-intensity matrix
Matrix of Byonic result after glycocylation selection

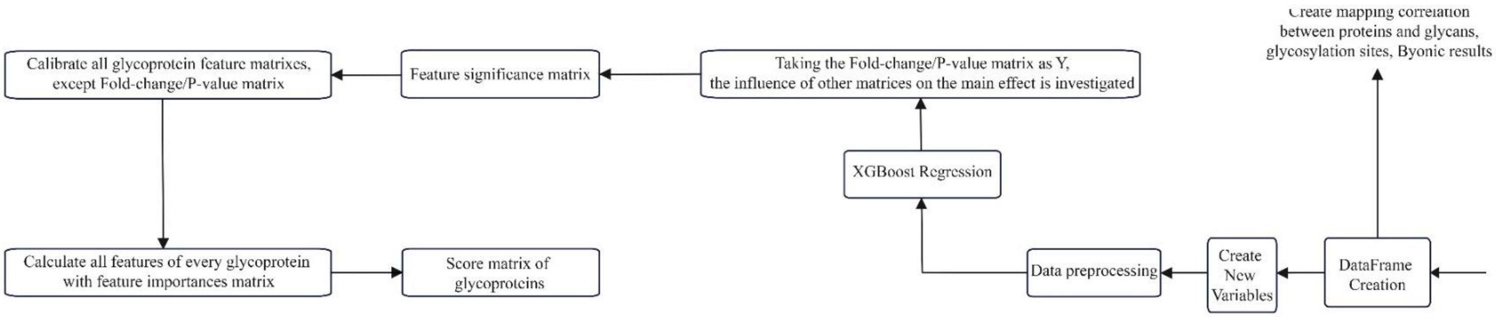
Create mapping correlation between proteins and glycans, glycosylation sites, Byonic results
DataFrame Creation
Create New Variables
Data preprocessing
XGBoost Regression
Taking the Fold-change/P-value matrix as Y, the influence of other matrices on the main effect is investigated
Feature significance matrix
Calibrate all glycoprotein feature matrixes, except Fold-change/P-value matrix
Calculate all features of every glycoprotein with feature importances matrix
Score matrix of glycoproteins

**Figure S5. Bootstrap resampling and effect size quantification.** Bootstrap is a statistical resampling technique that estimates properties like standard error, confidence intervals, and bias by generating multiple new samples from the original sample with replacement, allowing for the estimation of the statistic's distribution characteristics. This procedure further incorporates Cohen's d effect size to quantify the difference between the means of two groups relative to the standard deviation, which is used to correct the statistics of the resulting resampled set. In GDAS, the function of the Bootstrap method is to analyze the difference in sites or glycans between two groups, serving as a key step in feature engineering for data pre-processing. However, a limitation is the number of samples; if the count is less than 10, the analytical model becomes severely malformed, leading to a significant loss of analytical power with biological samples [19].

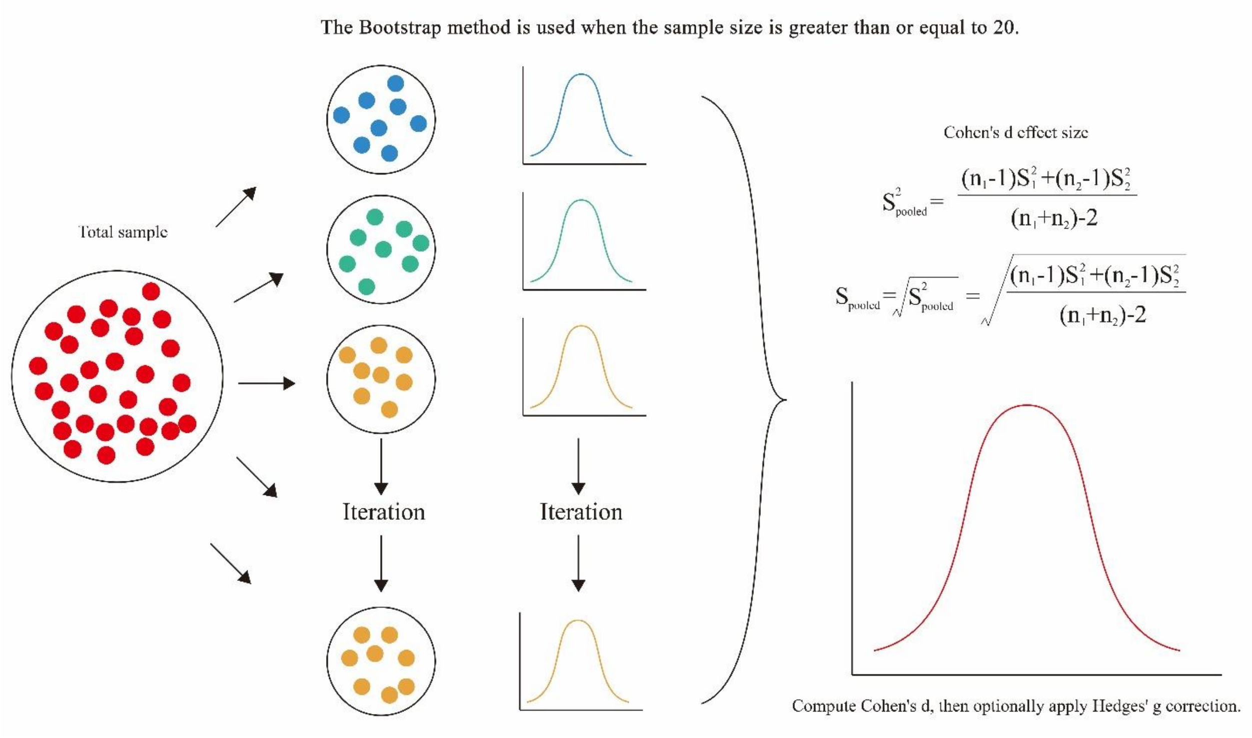

**Figure S6. Bayesian statistical inference.** Bayesian statistical inference, or Bayesian statistics, is a framework based on Bayes' theorem where a statistical distribution is iteratively fitted to a given prior distribution (an initial guess). Its core principle involves first establishing this prior distribution and then using the Markov-Chain Monte Carlo (MCMC) method to iterate until convergence, yielding a robust posterior distribution model of the sample data. As a supplementary method for feature engineering, Bayesian theory offers stronger analytical power, especially with fewer samples, with the core being the prior analytical model. In most biological prediction models, bivariate and univariate distributions are the most common statistical models used, though obtaining the maximum likelihood function often requires a substantial number of iterations [20].

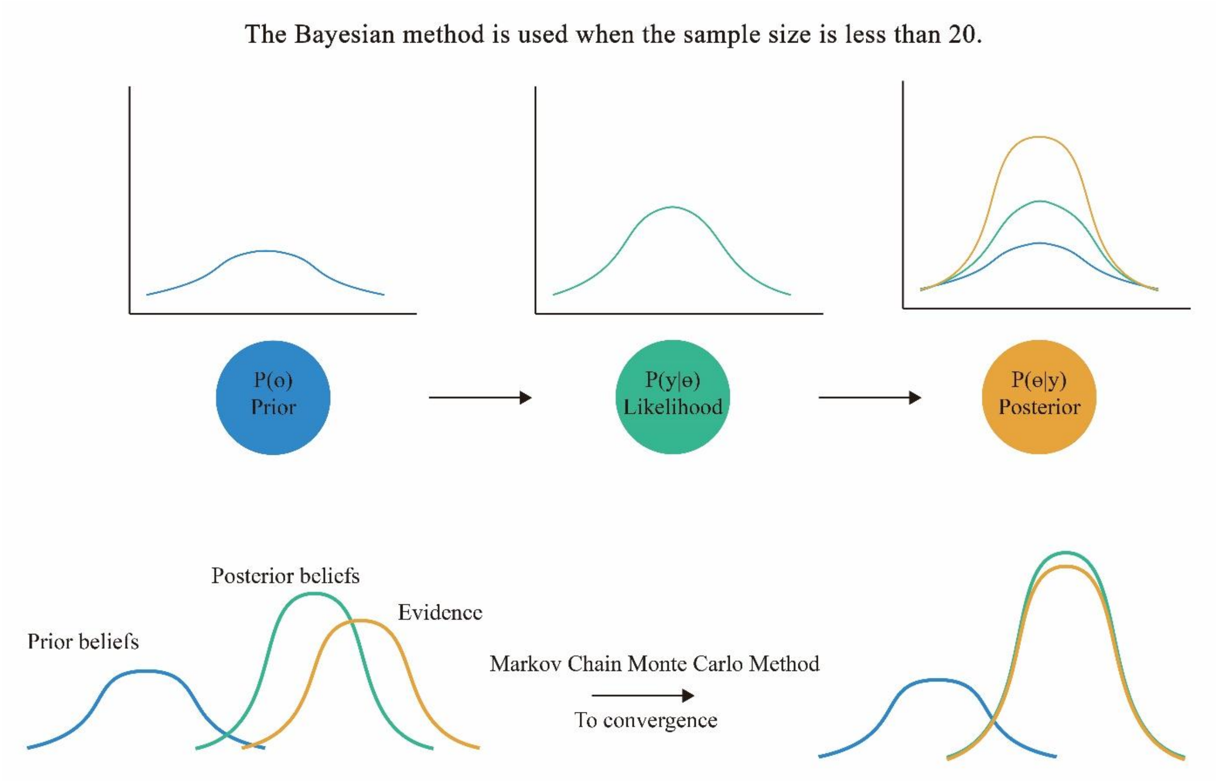

**Figure S7. Simplified diagram of the GDAS system.** The GDAS system is fundamentally divided into two loosely coupled parts: the Protein Screening and Analysis Module (comprising the Reload and Analysis Modules) and the Machine Learning Module (the Final Module). In the Protein Screening and Analysis Module, the intersection of detected proteins from two result sets is extracted and then filtered using user-defined thresholds to reduce the data volume. This module also performs statistical analysis, yielding data for meaningful visual evaluations, such as a volcano plot. The Machine Learning Module handles data reading, preprocessing, feature engineering, and integration of datasets from various sources, which are subsequently fed into the XGBoost algorithm for model fitting. The two modules are designed to minimize coupling, communicating only through data transmission, and are managed via a Graphical User Interface (GUI) that provides connection functions for the entire workflow.

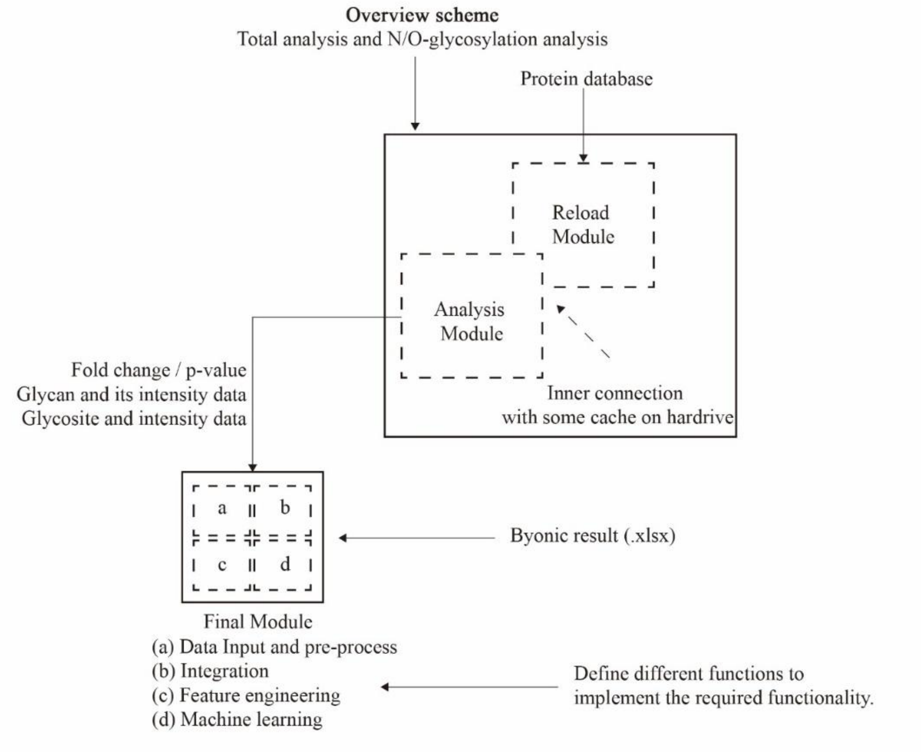

**Figure S8. eXtreme gradient boosting (XGBoost) algorithm.** XGBoost is a highly efficient and optimized machine learning algorithm that serves as an improved implementation of the Gradient Boosting Decision Tree (GBDT) framework. It constructs a powerful predictive model by sequentially integrating the predictions of multiple, simpler weak learners (decision trees). Furthermore, XGBoost is capable of handling data that contains many invalid values, which makes the algorithm exceptionally robust. By utilizing XGBoost as its core prediction algorithm, GDAS gains the ability to effectively discern the potential relationships between the experimental group and the control group based on the biological sample designation.

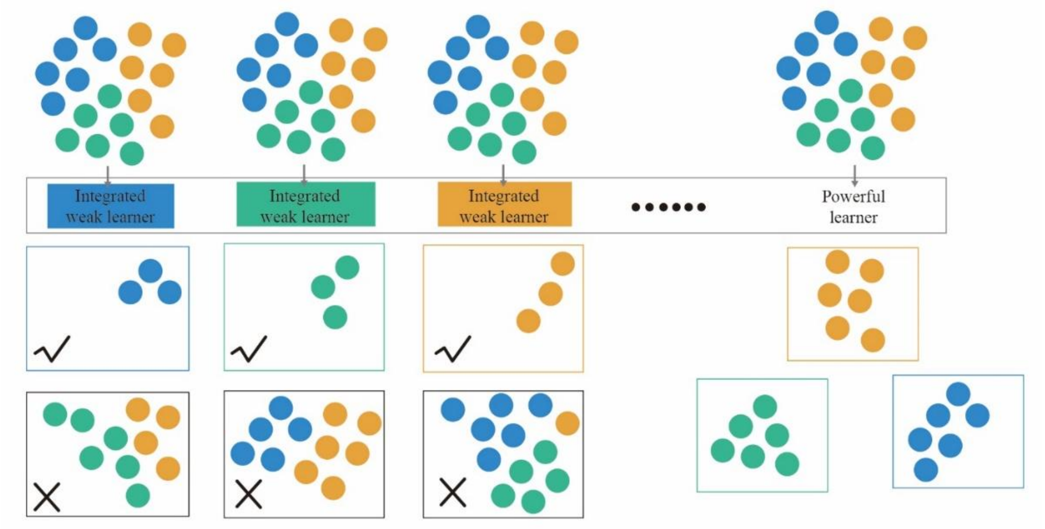

**Figure S9. GDAS MS/MS annotation of intact glycopeptides.** The GDAS program initially analyzes the raw mass spectrometry data and outputs key information for glycopeptides, including the presence of fucose, sialic acid, or high-mannose, and the Scan Number within the mass spectrometry file. This Scan Number is the core identifier used by the MSFileReader module to retrieve the corresponding retention time and spectral data, automatically generating a preliminary spectrum plot, along with an initial list of predicted glycans and peptides. To achieve the complete spectra annotation, this initial information is integrated with MSFileReader data, re-plotted in Adobe Illustrator (AI), where each key fragmentation peak is manually identified and labeled, and the comprehensive glycan information from GDAS is incorporated. The final, fully annotated spectrum image is then output from AI.

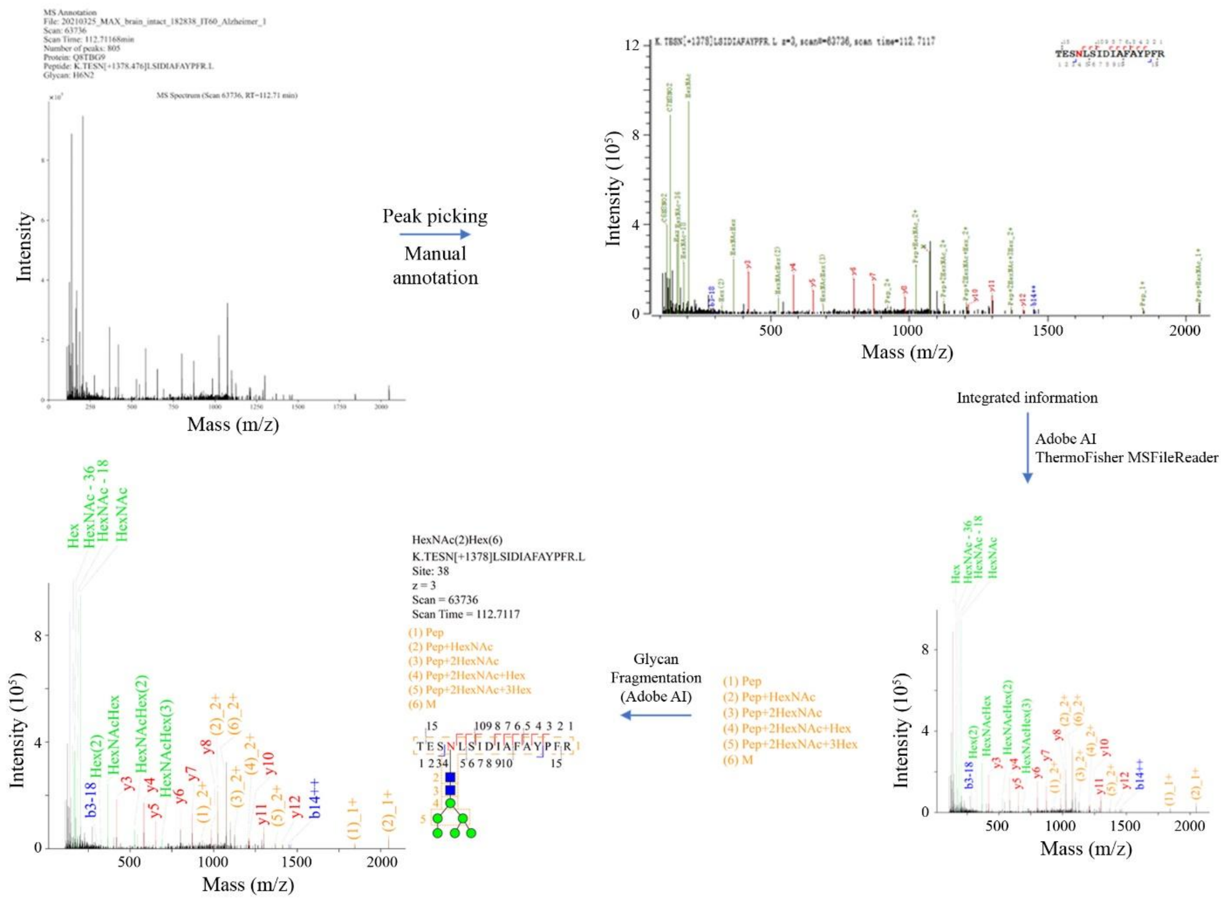
