## Supplementary material for "A Novel Glycoproteomics Platform for High-Throughput Identification of Disease-Associated Glycoforms": Installation guide

### GDAS Software Installation and User Guide

#### I. Installation Process

The installation involves both the core software and the deployment of Docker containers required for data processing.

1. Select installation directory: Launch the installer and browse to your preferred installation path.
2. Configure shortcuts: Choose whether to create a desktop shortcut for quick access.
3. File extraction: Click "Install" and wait for the installer to copy the necessary system files. This may take several minutes.
4. Docker image deployment: After the initial setup, a command window will automatically appear to deploy the required Docker images. Do not close this window until it finishes.
5. Complete setup: Once the deployment status indicates success, click "Finish" to exit the installer.

#### II. N-Glycosylation & O-Glycosylation Analysis

Use these modes to process glycopeptide data through the GDAS pipeline.

1. Initialize docker: Ensure the Docker Desktop application is running and that the GDAS images are listed in your Docker dashboard.
2. Launch GDAS: Open the GDAS Graphical User Interface (GUI).
3. Configure Resource allocation: Navigate to the menu bar and select Docker Threads. Enter the number of threads allocated for the container (the recommended default is 1).
4. Select processing mode: Choose either N-Glycosylation or O-Glycosylation based on your specific research requirements.
5. Define input paths: Input the directory paths for your required data files.
6. Set statistical thresholds: Enter your desired Fold-change and P-value cutoffs. Hover over the input fields to view the tooltip dialogue boxes for parameter guidance.
7. Execute: Click Run to begin the analysis.

**III. Final Prediction Mode (Step 2)**

This mode integrates previously processed data to generate finalized results.

1. Prerequisites: Verify that Docker is active and the GUI is open.
2. Thread configuration: Confirm that the Docker Threads setting is still correctly configured (usually 1).
3. Select mode: From the Modes dropdown menu, select Step 2 (Final Prediction).
4. Input file mapping: * Enter the file paths for the intermediate analysis files generated in the previous steps.
    - Click Run.
5. Select raw spectra: A tooltip prompt will appear; follow the instructions to select the corresponding raw spectrum files required to complete the prediction.
